# Single-cell RNA-seq reveals stepwise fate conversion of supporting cells to hair cells in the chick auditory epithelium

**DOI:** 10.1101/2022.08.25.505256

**Authors:** Mami Matsunaga, Ryosuke Yamamoto, Tomoko Kita, Hiroe Ohnishi, Norio Yamamoto, Takayuki Okano, Koichi Omori, Takayuki Nakagawa

## Abstract

In contrast to mammals, the avian cochlea, specifically the basilar papilla, can regenerate sensory hair cells, which involves fate conversion of supporting cells to hair cells. To determine the mechanisms for converting supporting cells to hair cells, we used single-cell RNA sequencing during hair cell regeneration in explant cultures of chick basilar papillae. We identified dynamic changes in the gene expression of supporting cells, and the pseudotime trajectory analysis demonstrated the stepwise fate conversion from supporting cells to hair cells. Initially, supporting cell identity was erased and transition to the precursor state occurred. A subsequent gain in hair cell identity progressed together with downregulation of precursor-state genes. Transforming growth factor beta receptor 1-mediated signaling was involved in induction of the initial step, and its inhibition resulted in suppression of hair cell regeneration. Our data provide new insights for understanding fate conversion from supporting cells to hair cells in avian basilar papillae.

## Introduction

Auditory hair cells (HCs) convert sound vibration into electrical potential that stimulates the auditory primary neurons and the loss of HCs diminishes auditory function. In the mammalian cochlea, virtually no HC regeneration occurs, leading to intractable sensorineural hearing loss. In contrast, in the avian cochlea, HCs are spontaneously restored after damage resulting in the maintenance of hearing throughout life. Therefore, the avian auditory sensory epithelium, known as the basilar papilla (BP), has been used as a model to study the mechanisms of HC regeneration for decades. Avian BP HCs are regenerated through two different pathways: direct conversion of supporting cells (SCs) to HCs (SC-to-HC conversion) and SC mitosis followed by differentiation into HCs (Stone and Cotanche 2007), with the former being the predominant mode of HC regeneration in explant cultures of chick BPs (Stone and Cotanche 2007; Shang et al., 2010). However, the detailed molecular mechanisms underlying HC regeneration in avian BPs have not been fully elucidated. To address this question, we established an explant culture model of chick BPs for HC regeneration and performed bulk RNA sequencing (RNA-seq) (Matsunaga et al., 2020). A notable characteristic of our explant culture model is that almost all new HCs were generated through SC-to-HC conversion (Matsunaga et al., 2020).

SC-to-HC conversion is also feasible in the mammalian cochlea by forced expression of Atoh1, a basic helix-loop-helix transcription factor that is required for differentiation of HCs from precursor cells during development (Bermingham et al., 1999; Chen et al., 2002; Woods et al., 2004). In the neonatal mouse cochlea, forced expression of Atoh1 by pharmacological inhibition of Notch signaling induces SC-to-HC conversion (Yamamoto et al., 2006; Doetzlhofer et al., 2009). Genetic manipulation to direct forced expression of Atoh1 induces SC-to-HC conversion in the mature mammalian cochlea (Kawamoto et al., 2003; Izumikawa et al., 2005; Kelly et al., 2012; Liu et al., 2012). However, its efficacy progressively decreases with age (Kelly et al., 2012; Liu et al., 2012). In addition, HCs generated by forced expression of Atoh1 exhibit the biophysical characteristics of HCs but fail to fully differentiate (Kelly et al., 2012; Liu et al., 2012; Liu et al., 2020), indicating that additional factors are required for the generation of mature HCs in adult mammals. More recently, the efficacy of additional factors in inducing SC-to-HC conversion has been reported in adult mice (Chen et al., 2021; Lee et al., 2020; Sun et al., 2021; Walters et al., 2017; Yamashita et al., 2018). However, to the best of our knowledge, sufficient recovery of auditory function has not yet been achieved in mammals.

In contrast to the mature mammalian cochleae, SCs in mature avian BPs can spontaneously upregulate Atoh1 after HC damage. In chick BPs, Atoh1 is highly expressed in SCs shortly after HC damage, and this upregulation occurs broadly across the BP (Cafaro et al., 2007). Not all Atoh1-expressing SCs show a change in cell fate to become new HCs (Cafaro et al., 2007). In addition, in developing mouse cochleae, many sensory precursor cells initially express Atoh1 and some Atoh1-expressing cells differentiate into HCs (Driver et al., 2013). However, the distinct mechanisms by which Atoh1-expressing SCs in chick BPs or precursor cells in developing mouse cochleae acquire the HC fate are unknown. A recent review of advancements and controversies in studies of stepwise fate conversion of several somatic cells to neurons indicates the importance of characterizing immature and intermediate phenotypes during direct conversion to facilitate the elucidation of a scientifically firm footing (Leaman et al. 2022). Thus, understanding the precise processes of SC-to-HC conversion in chick BPs may contribute to the achievement of more practical and sufficient SC-to-HC conversion in mature mammalian cochleae.

Molecular analyses of BP development have focused on major signaling pathways, including Notch (Daudet et al., 2005; 2009; Petrovic et al., 2014), fibroblast growth factor (Bermingham-McDonogh et al., 2001; Jacques et al., 2012), and Wnt signaling (Sienknecht and Fekete, 2008; Munnamalai et al., 2017). However, information on the gene expression patterns in developing or regenerating chick BPs in comparison to those in the zebrafish lateral line was limited (Lush et al., 2019; Baek et al., 2022). The baseline data of homeostatic BPs have been recently reported (Janesick et al., 2021) using single-cell RNA-seq, which provides fundamental information to analyze the changes in gene expression in SCs toward HC regeneration, similar to the findings in zebrafish (Lush et al., 2019; Baek et al., 2022) and mice (Kolla et al., 2020). To gain insights into the processes underlying SC-to-HC conversion in chick BPs, we conducted a high-resolution transcriptional analysis of SCs during SC-to-HC conversion in our explant culture model using single-cell RNA-seq.

## Results

### Hair cell regeneration occurs in explant cultures of chick basilar papillae

The BPs excised from post-hatch day 1 (P1) chicks were maintained for 24 h in the control media and provided explant cultures (Figure S1A). BPs consist of two types of sensory epithelial cells: HCs and SCs (Figure S1B). HCs are divided into two phenotypes: tall and short. Tall HCs are located in the neural portion of the BP and short HCs are present in the abneural portion (Figure S1B). Several types of non-sensory cells, including homogene cells, are present in the neural and abneural areas adjacent to sensory cells (Figure S1B). These non-sensory cells are connected to epithelial cells in the roof of the cochlear duct (Figure S1B).

Half of the BPs were exposed to streptomycin (SM), an ototoxic antibiotic, for 48 h to induce HC death, and the remaining BPs were maintained in control media for 48 h (Figure S1A). Subsequently, both samples were maintained in control media for an additional 96 h (Figure S1A). At the end of the culture period, the BPs were fixed and used for histological assessment. To assess the deletion of original HCs from cultured BPs and the generation of new HCs, we performed immunostaining for MYO7A and SOX2. We observed the two layers of the surface of the 40% area from the distal end of BPs (Figure S1C) and counted the number of HCs, which were labeled with MYO7A, and the newly generated, immature HCs, which were co-stained with MYO7A and SOX2. In samples that were cultured with SM, virtually no HCs were found in the 40% area following 48-h culture (day2_SM; Figure S1D), and a certain number of HCs were identified in the same region of BPs after additional 96-h culture in the control medium (day6_SM; Figure S1D). Immunohistochemistry demonstrated the presence of newly generated immature HCs in the specimens from day6_SM (Figure S1D). Quantitative assessments ascertained the total HC loss after 48 h of exposure to SM and the occurrence of HC regeneration (Figure S1F). These findings demonstrated that HC regeneration occurred during an additional 96 h of culture after 48 h of exposure to SM.

In BP cultures without SM exposure, HCs were well maintained during the 6-day culture period (Figure S1E). However, quantitative assessments indicated modest HC loss and the generation of new HCs during the 6-day culture, although the difference was not statistically significant among experimental groups (Figures S1G). These findings suggest that HC loss and subsequent regeneration also occurred in BP explants cultured in control media.

### Single-cell RNA sequencing of chick basilar papillae can identify clusters of supporting cells and neighboring non-sensory cells

To collect SCs at different stages of HC regeneration, we prepared five experimental groups of BP explant cultures: day0_pre (before explant culture), day2_SM (after 48-h SM exposure), day6_SM (96-h culture after 48-h SM exposure), day2_Ctrl (48-h culture without SM exposure), and day6_Ctrl (144-h culture without SM exposure) (Figure 1A). Histological assessments (Figure S1) indicated that these specimens contained intact SCs in the culture condition, SCs responding to HC injury at various stages of HC regeneration, and newly generated and intact mature HCs (Figure 1A). In addition to these five experimental groups, we collected BP samples that were immediately dissected from P1 chicks, namely, the intact group, which contained HCs and SCs *in vivo* (Figure 1A). Consequently, we harvested single cells from the six experimental groups (15-17 BPs per group).

**Figure 1.**
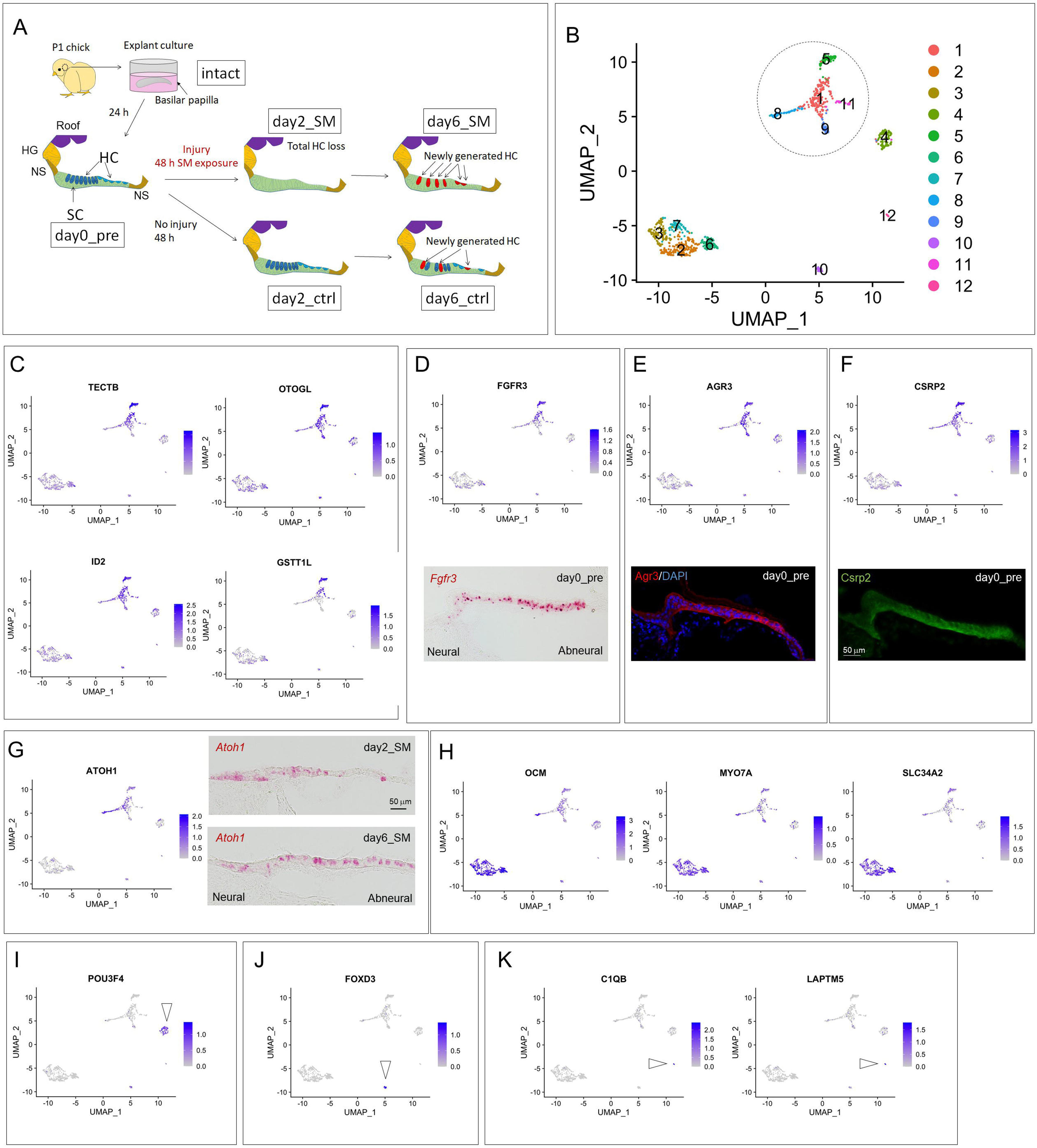
Single-cell RNA sequencing of chick basilar papillae identifies clusters of supporting cells and neighboring non-sensory cells. (A) Experimental groups. HC: hair cell, SC: supporting cell, HG: homogene cell, NS non-sensory cell, SM: streptomycin. (B) Uniform manifold approximation and projection (UMAP) plots of whole-cell populations. A circle indicates a group of clusters consisting of SCs and neighboring non-sensory cells. (C) UMAP plots for SC markers. (D-F) UMAP plots for FGFR3 (D), AGR3 (E), and CSRP2 (F) and histological distributions (*in situ* hybridization for *FGFR3*, immunostaining for AGR3 and CSRP2) in transverse sections of the basilar papillae (BPs) before exposure to streptomycin (day0_pre). Scale bar in (F) represents 50 μm for (D-F). (G) UMAP plots for ATOH1 and *in situ* hybridization for *ATOH1* in transverse sections of BPs after HC injury (day2_SM and day6_SM). Scale bar represents 50 μm. (H-K) UMAP plots for HC markers (H), a mesenchyme marker (I), a marker for neural crest-derived cells (J), and microglia markers (K). Arrowheads indicate corresponding clusters.

The capture of single cells dissociated from chick BPs and cDNA synthesis were performed using the C1 Single-Cell Auto Prep system (Fluidigm, San Francisco, CA, USA). Raw sequencing data were converted into FASTQ files using Illumina bcl2fastq software (Illumina, San Diego, CA, USA) and mapped to the chicken reference genome GRCg6a. After removing the cells with an RNA count of less than 20,000, detected genes less than 1,000, detected genes more than 4,000, or percentage of mitochondrial genes more than 12.5%, the remaining 1054 cells from chick BPs were normalized. Unsupervised clustering identified 12 clusters (Figure 1B).

The expression of known SC markers was examined to identify clusters containing SCs. Janesick et al. (2021) demonstrated the similarity of expressed genes in neighboring non-sensory cells to those in SCs (Janesick et al. 2021). Therefore, in this step, we extracted clusters containing SCs and neighboring non-sensory cells. The distributions of the SC marker genes *TECTB, OTOGL, ID2,* and *GSTTL1* indicated that a group of clusters 1, 5, 8, 9, and 11 included SCs (Figures 1B and 1C). We histologically examined the expression of *FGFR3*, AGR3, and CSRP2 as SC marker candidates, which were observed in SCs and neighboring non-sensory cells in day0_pre samples by *in situ* hybridization (ISH) or immunohistochemistry (Figures 1D-F). These markers were also found in clusters 1, 5, 8, 9, and 11 (Figures 1D-F). ATOH1, a key transcription factor for SC-to-HC conversion, is expressed in SCs in response to HC injury and converting HCs (Lewis et al., 2012). Histologically, *ATOH1* expression was broadly observed in SCs after HC injury by SM (Figure 1G), which is similar to previous observations (Cafaro et al., 2007; Lewis et al., 2012; Matsunaga et al., 2020). High expression of *ATOH1* was found in clusters 1 and 8 (Figure 1G), indicating that clusters 1 and 8 contained SCs responding to HC injury. Altogether, we identified clusters 1, 5, 8, 9, and 11 as SC clusters that contained SCs and neighboring non-sensory cells for subsequent analyses.

To characterize other clusters, we examined the distribution of HC markers (*OCM, MYO7A,* and *SLC34A2*; Janesick et al. 2021), mesenchymal cell markers (*POU3F4*; Coate et al., 2012; Brooks et al., 2019), neural crest-derived cell markers (*FOXD3*; Mundell and Labosky 2011; Lukoseviciute et al., 2018), and microglial markers (*LAPTM5* and *C1QB*; Bonham et al., 2019). All HC marker genes were strongly expressed in clusters 2, 3, 6, and 7 (Figure 1H). The expression of *POU3F4* was identified in cluster 4 (Figure 1I), and that of *FOXD3* was selectively detected in cluster 10 (Figure. 1J). Expressions of *LAPTM5* and *C1QB* were found in cluster 12 (Figure. 1K).

### Identification of supporting cells at different stages toward hair cell regeneration

We extracted clusters 1, 5, 8, 9, and 11 (Figure 1B), which contained SCs and neighboring non-sensory cells, and re-clustered this dataset. Eight SC clusters were obtained (Figures 2A and S2). Figure 2B shows the distribution of the SC clusters in each experimental group. The cells derived from the intact group were clearly separated from those harvested from the cultured BP samples, which were distributed in SC cluster 1 (Figure. 2A). Thus, we annotated SC cluster 1 as SCs *in vivo*.

**Figure 2.**
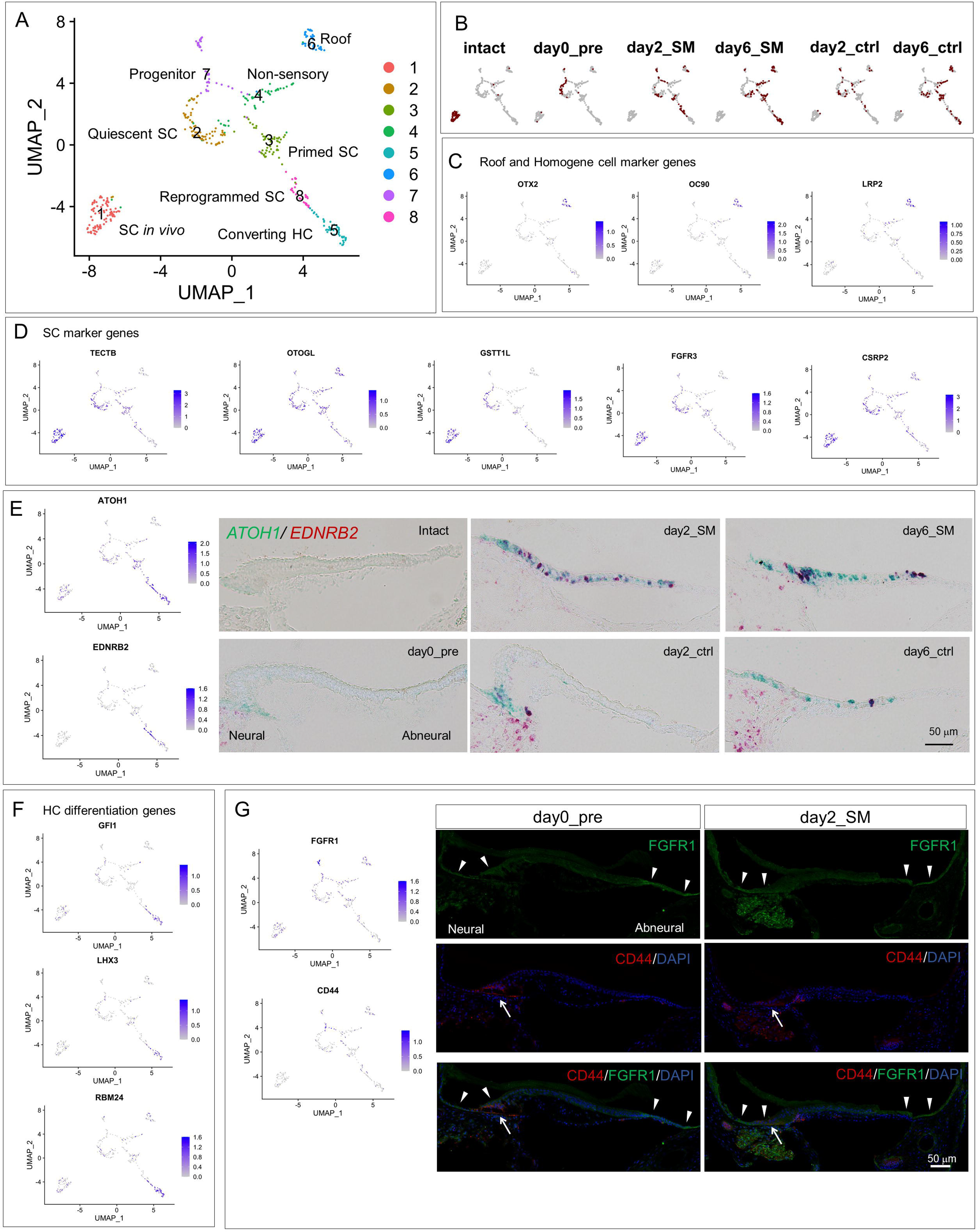
Identification of supporting cell clusters at different stages toward hair cell regeneration. (A) Uniform manifold approximation and projection (UMAP) plots of supporting cells and non-sensory cells. (B) UMAP plots for each experimental group. (C, D, F) UMAP plots for roof and homogene cell markers (C), supporting cell markers (D), and hair cell differentiation genes (F). (E) UMAP plots for ATOH1 and EDNRB2 and *in situ* hybridization for *ATOH1* (green) and *EDNRB2* (red) in transverse sections of basilar papillae (BPs) of each experimental group. Scale bar represents 50 μm. (G) UMAP plots for FGFR1 and CD44 and immunostaining for FGFR1 (green) and CD44 (red) in transverse sections of BP of day0_pre and day2_SM specimens. Arrowheads indicate FGFR1 expression, and arrows indicate CD44 expression. Scale bar represents 50 μm.

In the hierarchical cluster analysis, SC cluster 6 was clearly separated from the other SC clusters (Figure S2). SC cluster 6 was characterized by the expression of *OTX2*, *OC90,* and *LRP2* (Figures 2C, S3F, and Table S1). *OTX2* is a marker for the tegmentum vasculosum of the chick cochlear duct roof (Sanchez-Caderon et al., 2004), and *OC90* is expressed in the roof of the developing mouse cochleae (Hartman et al., 2015). Thus, SC cluster 6 has the characteristics of roof cells. *LRP2* is a specific marker for homogene cells (Janesick et al. 2021; 2022), and its expression is observed in the apical compartments of homogene cells adjacent to the roof of the cochlear duct (Janesick et al. 2021). Altogether, SC cluster 6 may be composed of apical compartments of homogene cells and roof cells. On the basis of these findings, we annotated SC cluster 6 as roof cells (Figure 2A).

SC cluster 4 also exhibited *LRP2* expression (Figure 2C), indicating that SC cluster 4 shares the identity of homogene cells. Hierarchical cluster analysis showed similarities among SC clusters 2, 3, and 4 (Figure S2). Among the three clusters, SC cluster 4 was characterized by low expression of SC marker genes (Figure 2D). In comparison with SC cluster 2 or 3, SC cluster 4 exhibited lower expression of several SC marker genes (*FGFR3, TECTB, GSTT1L,* and *LCAT*) (Figures S4A, S4B, and Table S3). On the other hand, in homeostatic chick BPs, Janesick et al. demonstrated the expression of SC marker genes including *TECTB, ID2,* and *TIMP3,* in the basal component of homogene cells and non-sensory cells adjacent to the sensory epithelium (Janesick et al. 2021). *CYR61* was included in the differentially expressed genes (DEGs) for SC cluster 4 (Figure S3D and Table S1), and its expression was observed in non-sensory cells adjacent to the sensory epithelium (Matsunaga et al., 2020). On the basis of these findings, SC cluster 4 was annotated as a cluster of non-sensory cells (Figure 2A).

Intense expression of SC marker genes (Figure 2D) and faint or virtually no expression of *LRP2* (Figure 2C) were characteristic of SC cluster 2. We then annotated SC cluster 2 as a quiescent SC. SC cluster 3 also showed expression of SC marker genes (Figure 2D), but in comparison with SC cluster 2, SC cluster 3 showed downregulation of several SC marker genes *(CSRP2, TECTB, FGFR3, LCAT, GSTT1L,* and *AGR3*) (Figure 3A and Table S2), indicating that while SC identity was still maintained in SC cluster 3, the erasure of SC identity had been initiated. In comparison with SC cluster 2, SC cluster 3 showed upregulation of *FGF10* (Figure 3A and Table S2). In addition, *SOX21* was included in the DEGs of SC cluster 3 (Figure S3C and Table S1). These genes are expressed by precursor cells in the embryonic chick BP (Freeman and Daudet 2012; Sanchez-Guardado et al., 2013), suggesting that the characteristics of SC cluster 3 are closer to those of precursor cells than SC cluster 2. Altogether, we annotated SC cluster 3 as primed SCs (Figure 2A).

**Figure 3.**
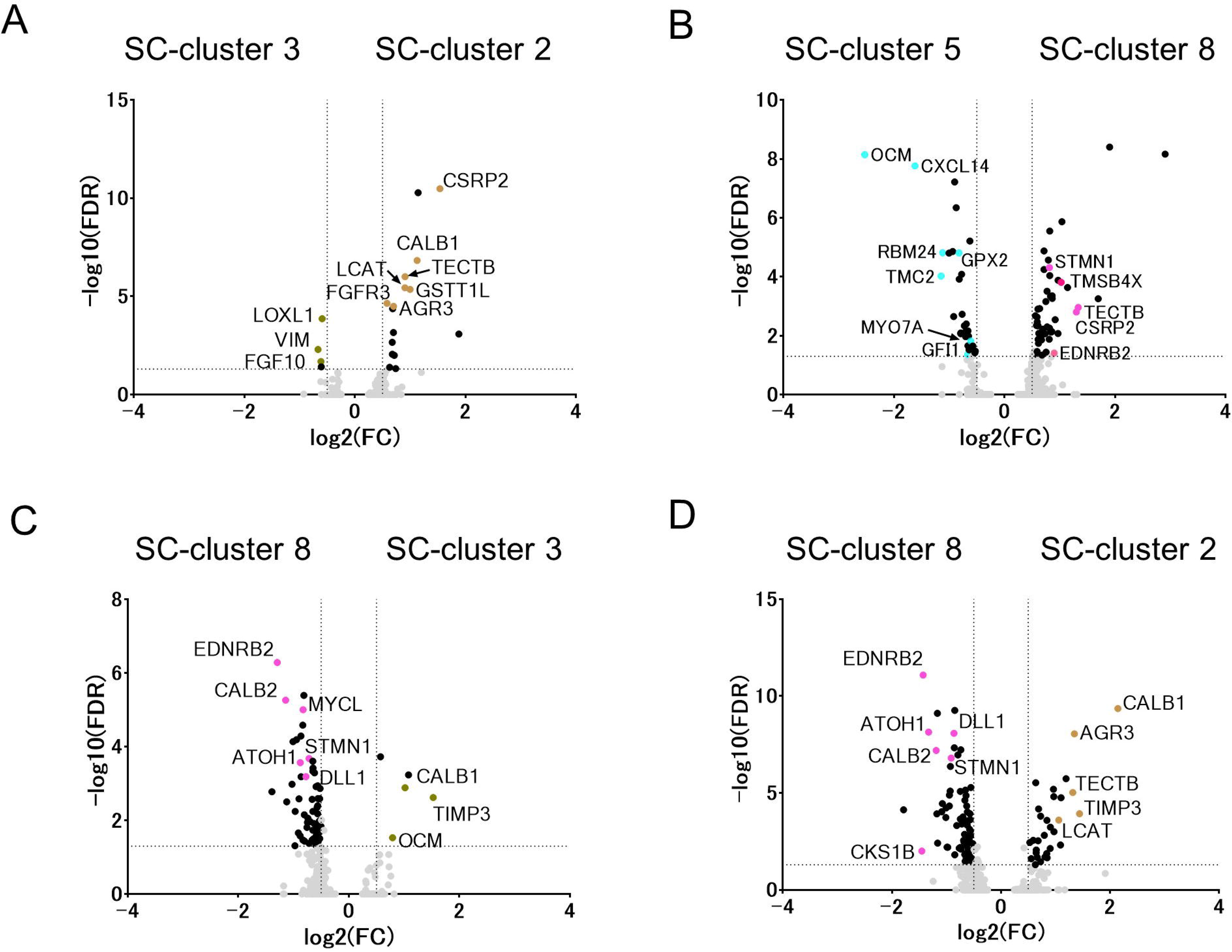
Volcano plots showing highly expressed genes (FDR < 0.05, log-fold-change threshold > 0.5) in comparisons between SC clusters 3 and 2 (A), SC clusters 5 and 8 (B), SC clusters 8 and 3 (C), and SC clusters 8 and 2 (D). Genes of interest are marked by colors corresponding to Figure 2A.

We highlighted the distribution of *ATOH1* expression in SC clusters. High expression of *ATOH1* was observed in SC clusters 5 and 8 (Figure 2E), and *ATOH1* was included in the DEGs of SC clusters 5 and 8 (Figures S3E and S3H, and Table S1). In SC cluster 5, high expression of the HC differentiation genes *GFI1* (Matern et al., 2020), *LHX3* (Hertzano et al., 2007), and *RBM24* (Cheng et al., 2020; Wang et al., 2021) was also identified (Figure 2F). In comparison with SC cluster 8, SC cluster 5 showed upregulation of HC marker genes (*MYO7A, GPX2, CXCL14, TMC2,* and *OCM*) and HC differentiation genes (*GFI1* and *RBM24*) (Figure 3B and Table S2). In contrast, SC cluster 8 exhibited higher expression of SC marker genes, including *TMSB4X, TECTB,* and *CSRP2*, than SC cluster 5 (Figure 3B and Table S2). These findings indicate that SC cluster 5 was in a more advanced stage of differentiation into HCs than SC cluster 8. Thus, we annotated SC cluster 5 as a cluster for converting HCs.

Compared to SC cluster 2 or 3, SC cluster 8 showed downregulation of SC marker genes (*AGR3, TECTB, LCAT, OCM,* and *TIMP3*) (Figures 3C, 3D, and Table S2), indicating the progression of erasure of SC identity in comparison with SC clusters 2 and 3. In addition, *DLL1* was identified in the DEGs of SC cluster 8 (Figure S3H and Table S1). In comparison with SC clusters 2 or 3, *DLL1* and *ATOH1* were upregulated in SC cluster 8 (Figures 3B, 3C, and Table S2), indicating that SC cluster 8 cells were directed to differentiate into HCs (Petrovic et al., 2014). Another characteristic of SC cluster 8 was the upregulation of the cell cycle-associated genes *MCM6, STMN1, CKS1B*, and *CDKN1C* (Figure S3H and Table S1). Wnt-associated genes *FZD9* and *SFRP2* (Figure S3H and Table S1), which are expressed in the precursor cells of chick BPs during development (Sienknecht and Fekete 2008), were also upregulated in SC cluster 8. Altogether, SC cluster 8 was directed to erase SC identity and to reverse cell identity to a developmentally immature status. Therefore, we annotated SC cluster 8 as reprogrammed SCs (Figure. 2A). The expression of *EDNRB2* was noted among the DEGs in SC cluster 8 (Figure S3H and Table S1). Furthermore, *EDNRB2* distribution was comparatively specific for SC cluster 8 (Figure 2E). We histologically examined the expression of *EDNRB2* in BPs together with *ATOH1*. In specimens from day2_SM, day6_SM, and day6_ctrl, some *ATOH1*-expressing SCs co-expressed *EDNRB2* (Figure 2E). On the basis of these results, we considered that *EDNRB2* could be a marker for reprogrammed SCs.

SC cluster 7 showed unique DEGs, including *PMP22, GNL3,* and *CD44* (Figure S3G and Table S1), which are known to play roles in the maintenance of progenitor or stem cell pools (Cai et al., 2017; Hagedorn et al., 1999; Tsai 2014; Tin et al., 2014; Chen et al., 2018; Zhu et al., 2020). In addition, *FGFR1* expression was noted in SC cluster 7 (Figures 2G, S3G, S4C, S4D, and Table S1). FGFR1 is crucial for the generation of the progenitor pool in the auditory sensory epithelium during murine cochlear development (Hayashi et al., 2010; Pirovola et al., 2002; Yang et al., 2019). On the basis of these findings, SC cluster 7 was annotated as a cluster of progenitor pools. To examine the localization of SC cluster 7 cells, immunohistochemistry for FGFR1 and CD44 was performed using specimens from day0_pre and day2_SM. FGFR1 expression was predominantly observed in non-sensory cells adjacent to the neural or abneural edges of the sensory epithelium (Figure 2G). CD44 expression was observed in non-sensory cells and SCs at the neural edge of the sensory epithelium (Figure 2G). Some CD44-positive cells co-expressed FGFR1 (Figure 2G). These findings indicate that SC cluster 7 cells may be present in the far neural and abneural regions of BPs, which is almost identical to the previously reported localization of stem cell-like populations in chick BPs (Janesick and Heller, 2019).

Consequently, SC clusters 2, 3, 5, and 8 were extracted as clusters containing SCs at different stages of SC-to-HC conversion for a subsequent pseudotime trajectory analysis.

### Pseudotime trajectory analysis reveals the stepwise fate conversion of supporting cells to hair cells

To characterize the processes underlying SC-to-HC conversion, we extracted the dataset of SC clusters 2, 3, 5, and 8 and performed a pseudotime trajectory analysis using Slingshot v1.6.0 (Street et al., 2018). The uniform manifold approximation and projection (UMAP) plots exhibited an approximately linear distribution of SCs (Figure 4A). We examined alterations in the expression levels of the genes of interest along a pseudotime trajectory. To classify genes according to alteration patterns along a pseudotime trajectory, we identified 500 DEGs along a pseudotime trajectory by fitting a random forest regression model (Wright and Ziegler, 2019). The top 500 genes that showed alterations along a pseudotime were extracted and classified into six time-series clusters (Figures 4D-I, S5, and Table S4). In Figures 4B-I and S5, the x-axis represents the pseudotime (0-1.0) and the y-axis shows the normalized expression value of the genes of interest.

**Figure 4.**
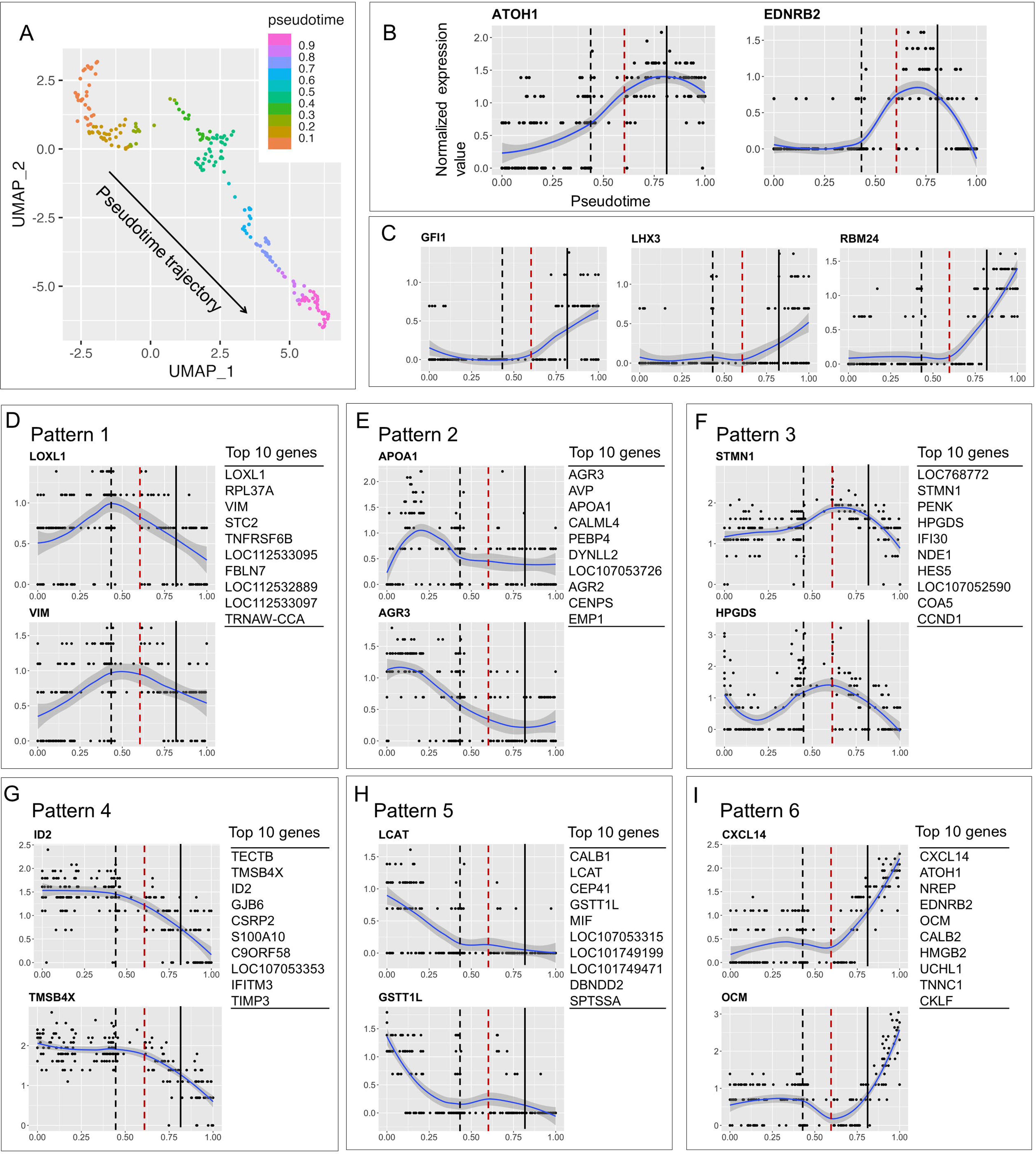
Pseudotime trajectory analysis reveals the stepwise fate conversion of supporting cells to hair cells. (A) Uniform manifold approximation and projection (UMAP) plots of the SC clusters 2, 3, 5, and 8 in Figure 2A. A pseudotime ordering was inferred by Slingshot v1.6.0. (B, C) Normalized expression values of *ATOH1* and *EDNRB2* (B) and hair cell differentiation genes (C) along a pseudotime line. Black dotted, red dotted, and black lines indicate the timing of *EDNRB2* induction, induction of hair cell differentiation genes, and *ATOH1* downregulation, respectively. (D-I) Normalized expression values of representative genes for six time-series clusters (patterns 1-6) along a pseudotime line and the top 10 genes for each cluster.

First, we focused on alterations in the expression of *ATOH1* and *EDNRB2* (Figure 4B). These genes were included in pattern 6 of the time-series clusters. *ATOH1* upregulation was initiated from time point 0 on the pseudotime, and its peak was observed at approximately 0.8 (Figure 4B). The peak of *ATOH1* expression is marked by a black line (Figures 4B-I and S5), which indicates the time point for the initiation of *ATOH1* downregulation. *EDNRB2* upregulation was initiated at around 0.4 on the pseudotime (black dotted line in Figures 4B-I and S5) and reached the peak slightly earlier than *ATOH1* (Figure 4B). To determine the timing of the initiation of HC differentiation, we examined the expression patterns of HC differentiation genes, including *GFI1, LHX3,* and *RBM24* (Figure 4C). The upregulation of these genes was initiated at around 0.6 on the pseudotime trajectory (red dotted lines in Figures 4B-I and S5), slightly later than *EDNRB2* induction.

Figures 4D-I show the representative alterations and the top 10 genes for each time-series cluster. Patterns 1-3 and 6 showed transient or continuous upregulation, and patterns 4 and 5 exhibited downregulation along a pseudotime. Pattern 1 exhibited transient upregulation up to *EDNRB2* induction (Figure 4D). Pattern 2 showed transient upregulation before *EDNRB2* induction (Figure 4E). The peaks of transient upregulation of pattern 3 genes appeared around the initiation of HC differentiation gene induction (Figure 4F). Pattern 6 included both transiently and continuously upregulated genes, and peaks of transiently upregulated genes were observed after the induction of HC differentiation genes (Figures 4B, 4I, and S5F). The continuously upregulated genes included HC marker (*CXCL14* and *OCM*, Figure 4I) and nascent HC marker (*NREP* and *CALB2*, Figure S5F) genes (Janesick et al., 2022). The transiently upregulated genes included *ATOH1, EDNRB2*, *HMBG2,* and *UCHL1* (Figures 4B and S5F), which are expressed by progenitor cells in various organs (Abraham et al., 2013; Bondurand et al., 2018; Driver et al., 2013; Sakurai et al., 2006). The induction of HC marker genes was initiated at the same time as HC differentiation gene induction (Figure 4I), while the other genes in pattern 6, including nascent HC marker genes, were upregulated after *EDNRB2* induction (Figure S5F). Regarding the downregulated genes, pattern 4 genes appeared downregulated after the *EDNRB2* induction (Figure 4G), whereas pattern 5 genes were downregulated before the *EDNRB2* induction (Figure 4H). Many SC marker genes were included in patterns 4 (*TECTB, TMSB4X, ID2, CSRP2,* and *TIMP3*; Figs. 4G and S5D) and 5 (*LCAT* and *GSTT1L*; Figure 4H).

From the point of view of SC and HC identities, the expression levels of SC marker genes were almost maintained and those of HC differentiation genes were not upregulated before *EDNRB2* induction (black dotted lines in Figures 4 and S5). In contrast, the expression level of *ATOH1* gradually increased before *EDNRB2* induction. Thus, we annotated this period (before *EDNRB2* induction) as the priming stage for SC-to-HC conversion. After *EDNRB2* induction, most SC marker genes were downregulated (Figures 4G and S5D). Slightly later, upregulation of HC differentiation genes was initiated (Figure 4C). During the period between *EDNRB2* induction and HC differentiation gene induction, the erasure of SC identity began, but the acquisition of HC identity was not initiated. Therefore, this period was annotated as the initial stage. After HC differentiation gene induction, the erasure of SC identity and acquisition of HC identity progressed simultaneously. The period between HC differentiation gene induction and *ATOH1* downregulation was annotated as the intermediate stage. SCs in this period exhibited high expression of progenitor marker genes, including pattern 6 genes, suggesting that SCs were reprogrammed to an immature state during this period. After *ATOH1* downregulation, which is a sign of initiation for the maturation process of regenerated HCs (Cafaro et al., 2007), the acquisition of HC identity progressed further, and progenitor marker genes were downregulated. This stage was annotated as the late stage. Thus, HC specification may occur during the intermediate stage.

### Transforming growth factor beta signaling is involved in the initiation of fate conversion of supporting cells to hair cells

As candidates for triggers for *EDNRB2* induction, we focused on pattern 1 genes, which were upregulated up to *EDNRB2* induction (Figure 4D). Among the pattern 1 genes, we highlighted *LOXL1* and *VIM*, because these two genes exhibited remarkable upregulation in pattern 1 genes and are associated with epithelial–mesenchymal transition (EMT) (Cheng et al., 2016; Hu et al., 2020). EMT is involved in the induction of regenerative processes (Oh et al., 2018; Marconi et al., 2021) and the gain of epithelial stem cell properties (Mani et al., 2008). Mouse utricle epithelial cells also acquire the features of prosensory cells via EMT (Zhang and Hu, 2012). Therefore, we focused on transforming growth factor beta (TGFb) signaling as upstream of both LOXL1(Voloshenyuk et al., 2011; Xie et al., 2013; Ma et al., 2018; Ma et al., 2021) and VIM (Cheng et al., 2016; Vervoort et al., 2013). In addition, TGFb1 application induced upregulation of *LOXL1* and downregulation of *LCAT* in mouse astrocyte primary cultures (Hamby et al., 2012), which is consistent with the present findings before *EDNRB2* induction in a pseudotime trajectory analysis (Figures 4D and 4H). The expression of *TGFBR1* was observed in SC clusters 2, 3, and 8 (Figure S6). Thus, we examined the effects of pharmacological inhibition of TGFb receptor1 (TGFbR1) on the transition from the priming stage to the initial or intermediate stage by assessing *EDNRB2* expression.

Chick BP explants were cultured with SM alone or SM and the TGFbR1 inhibitor SB505124 (Byfield et al., 2004), which is a selective inhibitor of TGFbR1 (ALK4, ALK5, and ALK7), at a concentration of 2 μM for 48 h (Figure 5A). In both BPs cultured with SM alone and SM and SB505124, total HC loss was confirmed. Quantitative real-time polymerase chain reaction (qPCR) demonstrated a significant increase in *EDNRB2* expression in samples incubated with SM alone compared to those before SM exposure (Figure 5B). Supplementation with SB505124 significantly reduced *EDNRB2* expression 48 h after SM exposure (Figure 5B). Thus, 48-h exposure to SM induced *EDNRB2* upregulation in BP explant cultures, and pharmacological inhibition of TGFbR1 suppressed its expression. We also examined the histological expression of *EDNRB2*. ISH for *EDNRB2* demonstrated a trend of decreasing *EDNRB2*–expressing SCs in BPs following supplementation with SB505124 (Figure 5C). We quantified *EDNRB2*–expressing cells by counting nuclei stained with 4′,6-diamidino-2-phenylindole (DAPI) in *EDNRB2*–expressing cells in three transverse sections of BPs from the distal (0%-30% from the distal end), mid (30%-70%), and proximal (70%-the proximal end) portions of each sample. The numbers of *EDNRB2*–expressing cells in BPs treated with SM and SB505124 were significantly lower than those in BPs treated with SM alone (Figure 5C), which is compatible with the results of qPCR (Figure 5B). Thus, inhibition of TGFbR1 suppressed *EDNRB2* expression in BP explants after HC injury.

**Figure 5.**
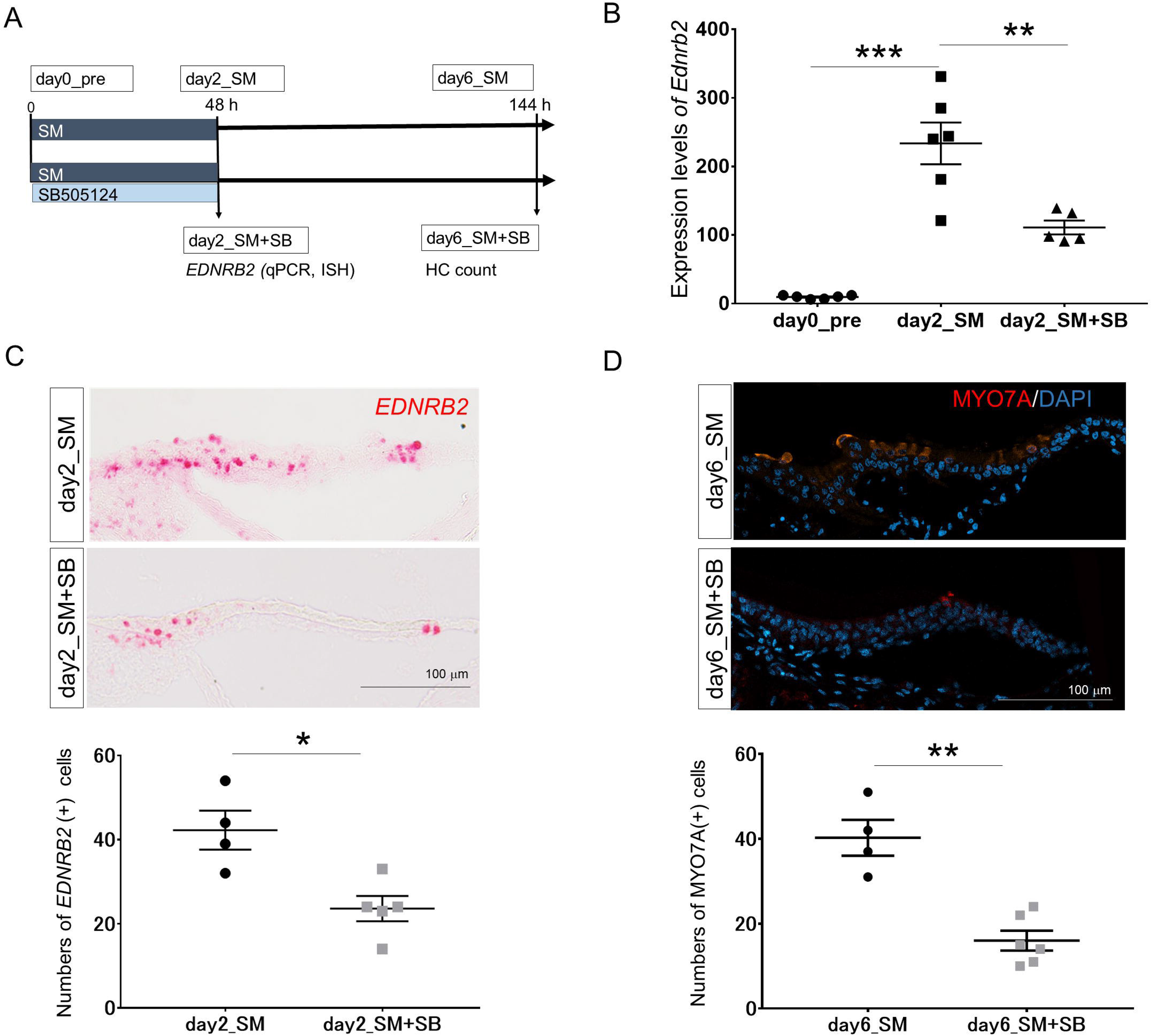
Pharmacological inhibition of transforming growth factor β type 1 receptor suppresses fate conversion of supporting cells to hair cells. (A) Experimental design and groups. SM: streptomycin, HC: hair cell, qPCR: quantitative real-time polymerase chain reaction, ISH: *in situ* hybridization. (B) Quantitative PCR reveals a significant increase in *EDNRB2* expression levels by 48-h exposure to SM (day0_pre [n=6] vs. day2_SM [n=6]) and significant suppression by the TGFbR1 inhibitor SB505124 (day2_SM [n=6] vs. day2_SM+SB [n=5]). Error bars represent standard errors. Welch t-test was used to calculate *P* values. \*\**P* < 0.01, \*\*\**P* < 0.001. (C) *In situ* hybridization for *EDNRB2* (red) in transverse sections of the basilar papillae (BPs) in the day2_SM and day2_SM+SB samples. Scale bar represents 100 μm. The number of *EDNRB2-*expressing cells in day2_SM+SB samples (n=5) is significantly lower than that in day2_SM samples (n=4). Error bars represent standard errors. Welch t-test was used to calculate *P* values. \**P* < 0.05. (D) Regenerated hair cells are labeled by MYO7A (red) and nuclear staining with DAPI (blue) in transverse sections of the basilar papillae (BPs) in day6_SM (n=4) and day6_SM+SB samples (n=6). Scale bar represents 100 μm. The number of regenerated hair cells in day6_SM+SB samples was significantly lower than that in day6_SM samples. Error bars represent standard errors. Welch t-test was used to calculate *P* values. \*\**P* < 0.01.

Next, we examined the effects of *EDNRB2* suppression by a TGFbR1 inhibitor on the generation of new HCs in specimens cultured for an additional 96 h after 48 h of SM exposure (Figure 5A). The number of MYO7A-positive cells in specimens treated with SM and SB505124 (day6_SM+SB) was significantly lower than those treated with SM alone (day6_SM) (Figure 5D). Altogether, inhibition of TGFbR1 repressed the transition from the priming stage to the initial or intermediate stage of SC-to-HC conversion, resulting in a reduction in regenerated HCs.

## Discussion

### Single-cell RNA sequencing reveals stepwise fate conversion from supporting cells to hair cells

For single-cell RNA-seq of SCs at different stages of SC-to-HC conversion, dissociated single cells were harvested from chick BPs before and after explant cultures with or without HC damage. Unsupervised clustering of the whole-cell populations identified 12 clusters. We extracted five clusters containing SCs and neighboring non-sensory cells. Re-clustering of this subpopulation identified eight clusters. Based on the gene expression patterns and histological assessments, we extracted four clusters consisting of SCs at different stages of SC-to-HC conversion. A pseudotime trajectory analysis of the extracted SC clusters revealed the time courses of alterations in expressed genes during SC-to-HC conversion, which indicated the stepwise conversion of SCs to nascent HCs. We divided the SC-to-HC conversion processes into the priming, initial, intermediate, and late stages according to alterations in gene expression associated with SC or HC identity along a pseudotime trajectory (Figure 6). The priming stage was characterized by gradient upregulation of *ATOH1* and transient upregulation of several genes, including *LOXL1* and *VIM*. In the initial stage, erasure of SC identity, not acquisition of HC identity, was induced. The characteristic genes for the intermediate stage, which can be associated with reversal to a precursor cell state, and HC marker genes were upregulated in the intermediate stage. In the late stage, HC differentiation progressed in parallel with the downregulation of precursor genes. Similar to HC regeneration in gene-manipulated neonatal mouse cochleae (Li et al., 2022) and zebrafish (Baek et al., 2022), HC regeneration in chick BPs occurs in a stepwise fashion, and reversal to a precursor cell state may be a crucial process.

**Figure 6.**
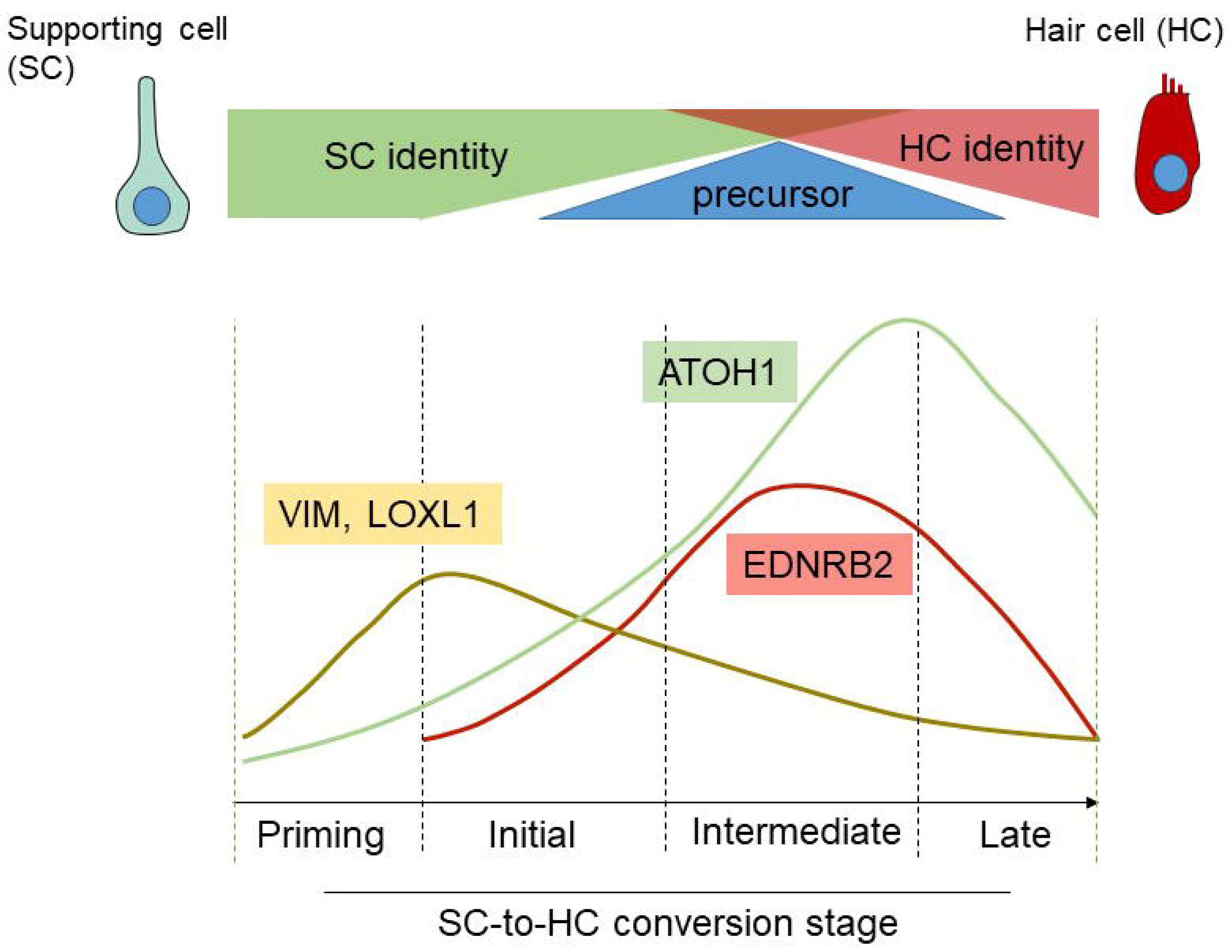
Schematic drawing of stepwise conversion from supporting cells to hair cells. Alterations in gene expression associated with supporting cell (SC) and hair cell (HC) identities along a pseudotime line indicate that SC-to-HC conversion occurs in four steps, i.e., the priming, initial, intermediate, and late stages. The priming stage is characterized by a gradual increase in ATOH1 and the upregulation of VIM and LOXL1. The initial stage is characterized by erasure of SC marker genes and induction of EDNRB2 upregulation. The intermediate stage is characterized by the initiation of HC identity acquisition and high expression of precursor-associated genes. The late stage is characterized by upregulation of HC marker genes and downregulation of precursor-associated genes.

### Initiation of fate conversion from supporting cells to hair cells

We focused on *LOXL1* and *VIM* as candidate switch genes for SC-to-HC conversion and highlighted TGFb signaling as a common upstream modulator of LOXL1 and VIM. A recent study focusing on LIN28B and FST indicated the involvement of TGFb signaling in the induction of SC reprogramming into precursor-like cells in the mouse cochlea (Li et al., 2022). The necessity of distinct control of TGFb signaling activity by FST for sufficient induction of SC reprogramming has also been demonstrated (Li et al., 2022). HC regeneration by the co-activation of LIN28B and FST was diminished by TGFb2, a TGFbR1 ligand, in explant cultures of postnatal mouse cochleae, suggesting that excessive activation of TGFbR1 represses HC regeneration (Li et al., 2022). In the present study, significant downregulation of *TGFB2* was identified in a cluster of converting HCs, SC cluster 5 (Figures S3E and S7A). In a pseudotime trajectory, downregulation of *TGFB2* was apparent in the late stage of SC-to-HC conversion (Figure S7B). In short, in our model, TGFbR1 may be activated in the priming stage of SC-to-HC conversion and thereafter suppressed during the process of HC differentiation in chick BP explants, which is compatible with the regulation of TGFb signaling during HC regeneration induced by the co-activation of LIN28B and FST in mouse cochleae (Li et al., 2022). In contrast, in the developing mouse cochleae, pharmacological inhibition of TGFbR1 suppresses the differentiation of precursor cells into outer HCs (Kolla et al., 2020). Altogether, TGFbR1-mediated signaling may be involved in both the induction of SC reprogramming and HC differentiation from precursor state cells. Therefore, we supplemented a TGFbR1 inhibitor to the culture medium only for the initial 48 h of the culture period to suppress TGFbR1-mediated signaling in the priming stage of SC-to-HC conversion. In the subsequent 4-day culture, BP explants were incubated with the control culture media to avoid the possible influences of a TGFbR1 inhibitor on HC differentiation in the later stage. The results showed that a TGFbR1 inhibitor significantly reduced *EDNRB2* expression in chick BPs damaged by SM and subsequent HC regeneration (Figure 5), indicating the involvement of TGFbR1-mediated signaling in the induction of SC reprogramming and the importance of this process for HC regeneration.

### Possible mechanisms for hair cell specification

The present results suggest that HC specification occurs during the intermediate stage of SC-to-HC conversion. Thus, highly expressed genes in the intermediate stage, namely, *EDNRB2*, *HMGB2,* and *UCHL1*, could contribute to the HC specification of reprogrammed SCs.

Mammals have two subtypes of endothelin receptors, EDNRA and EDNRB, while the chicken has three subtypes, EDNRA, EDNRB1, and EDNRB2 (Liu et al., 2019). EDNRB2 stimulates intracellular calcium, mitogen-activated protein kinase/extracellular signal-regulated kinase, and cAMP/protein kinase A signaling pathways, similar to mammalian EDNRB (Liu et al., 2019). EDNRB modulates the differentiation of Schwann cell precursors (Brennan et al., 2000; Quintes et al., 2016) and glial cell precursors (Hammond et al., 2015). *HMGB2* encodes high-mobility group box2, a chromatin-associated protein that regulates transcription, cell proliferation, and differentiation (Yanai et al., 2009). HMGB2 is involved in the adipogenesis of mesenchymal stem cells (Lee et al., 2018; Chen et al., 2021) and hematopoiesis in hematopoietic stem cells (Zhang et al., 2020). In addition, HMGB2 regulates the neurogenic-to-gliogenic fate transition of neural stem/progenitor cells via epigenetic modifications (Bronstein et al., 2017). *UCHL1* encodes ubiquitin C-terminal hydrolase L1, also known as PGP9.5, a deubiquitinating enzyme that is abundant in neurons (Wilkinson et al., 1989) and plays a crucial role in metabolic regulation (Alpaugh et al., 2021; Reinicke et al., 2019). UCHL1 enhances the differentiation of neural progenitor cells into neurons (Sakurai et al., 2006) and inhibits neurodegeneration by responding to energy requirements and endoplasmic reticulum stress (Reinicke et al., 2019). In addition, the nascent HC marker genes *NREP* and *CALB2* also initiate upregulation in the initial stage, similar to *EDNRB2*, and exhibit comparatively high expression levels in the intermediate stage (Figure S5F), suggesting the possible roles of these genes in HC specification. The distinct roles of the above-mentioned molecules in HC specification should be examined in future studies.

### Comparison with an *in vivo* hair cell regeneration model of chick basilar papillae

A single-cell RNA-seq analysis for the early phase of HC regeneration processes in chick BPs *in vivo* recently demonstrated distinct markers that are expressed in SCs responding to HC damage and in newly regenerated, nascent HCs (Janesick et al., 2022). In this *in vivo* model, HC regeneration occurred through both SC-to-HC conversion and SC mitosis, followed by differentiation into HCs (Janesick et al., 2022). In the dataset of this study, significant upregulation of *UCHL1, VIM, STMN1,* and *HPGDS* was identified in responding SCs in comparison with quiescent SCs (Janesick et al., 2022). In comparison with mature HCs, *NREP, CALB2, STMN1, UCHL1, HMGB2,* and *EDNRB2* are significantly upregulated in nascent HCs (Janesick et al., 2022). These findings are compatible with the results of a pseudotime trajectory analysis in the present study (Figures. 4, S5). The responding SCs of an *in vivo* model (Janesick et al., 2022) will correspond to SCs in the initial or early phase of the intermediate stage in our model, and nascent HCs of an *in vivo* model may correspond to SCs in the late phase of the intermediate stage or converting HCs in the late stage in our model. These genes are strong candidates for characterizing immature and intermediate phenotypes during the fate conversion of SCs to HCs in chick BPs.

In the present study, single-cell RNA-seq revealed the stepwise conversion from SCs to HCs through reversal to precursor status in chick BP explant cultures. *EDNRB2* is specifically expressed in SCs reprogrammed for HC regeneration. TGFb signaling is involved in the initiation of SC-to-HC conversion. Our data provide new insights into the mechanisms of SC-to-HC conversion in chick BPs and will contribute to the exploration of critical signaling pathways for HC regeneration through SC-to-HC conversion in future studies.

## Materials and Methods

### Key resource table

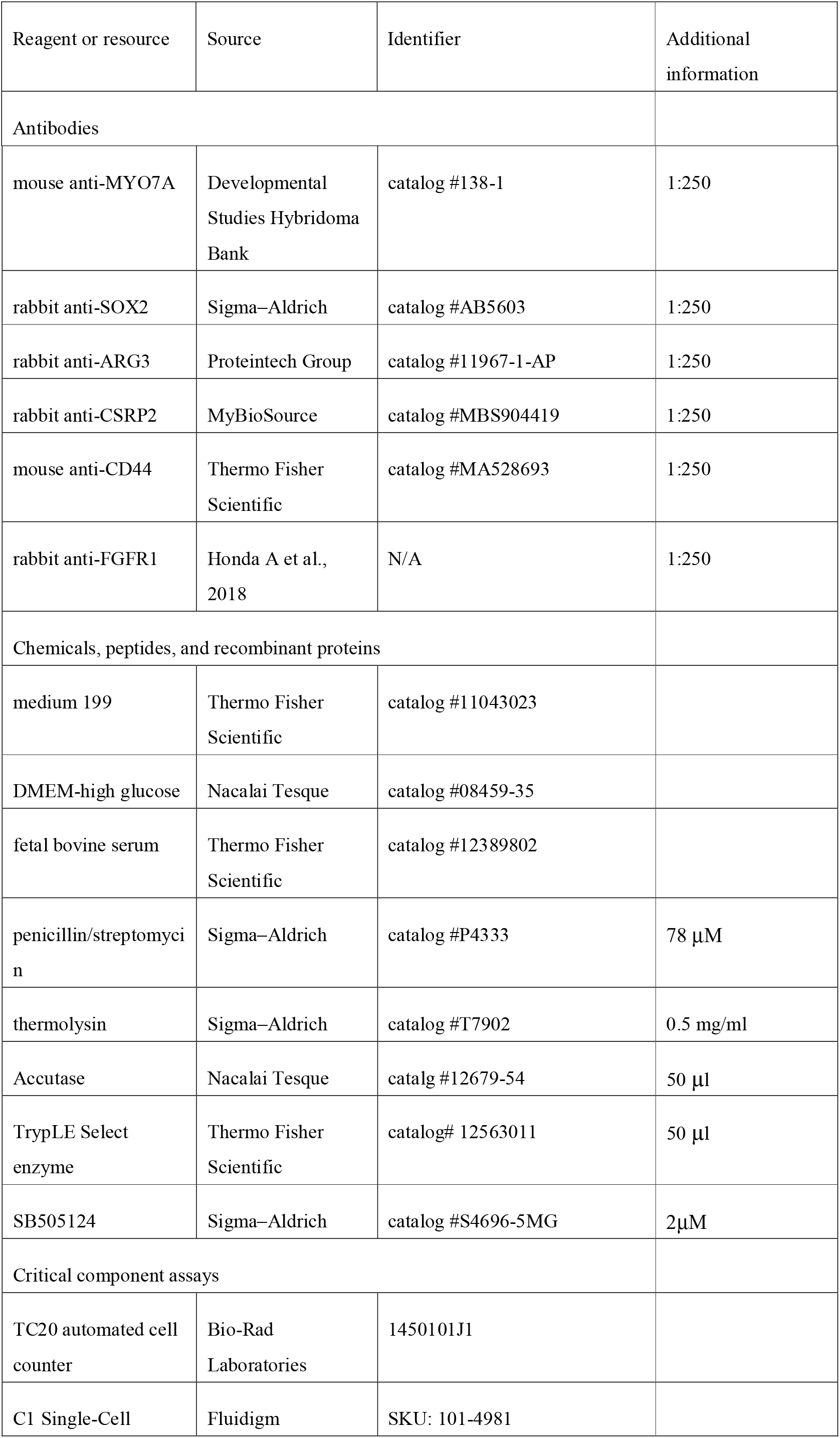
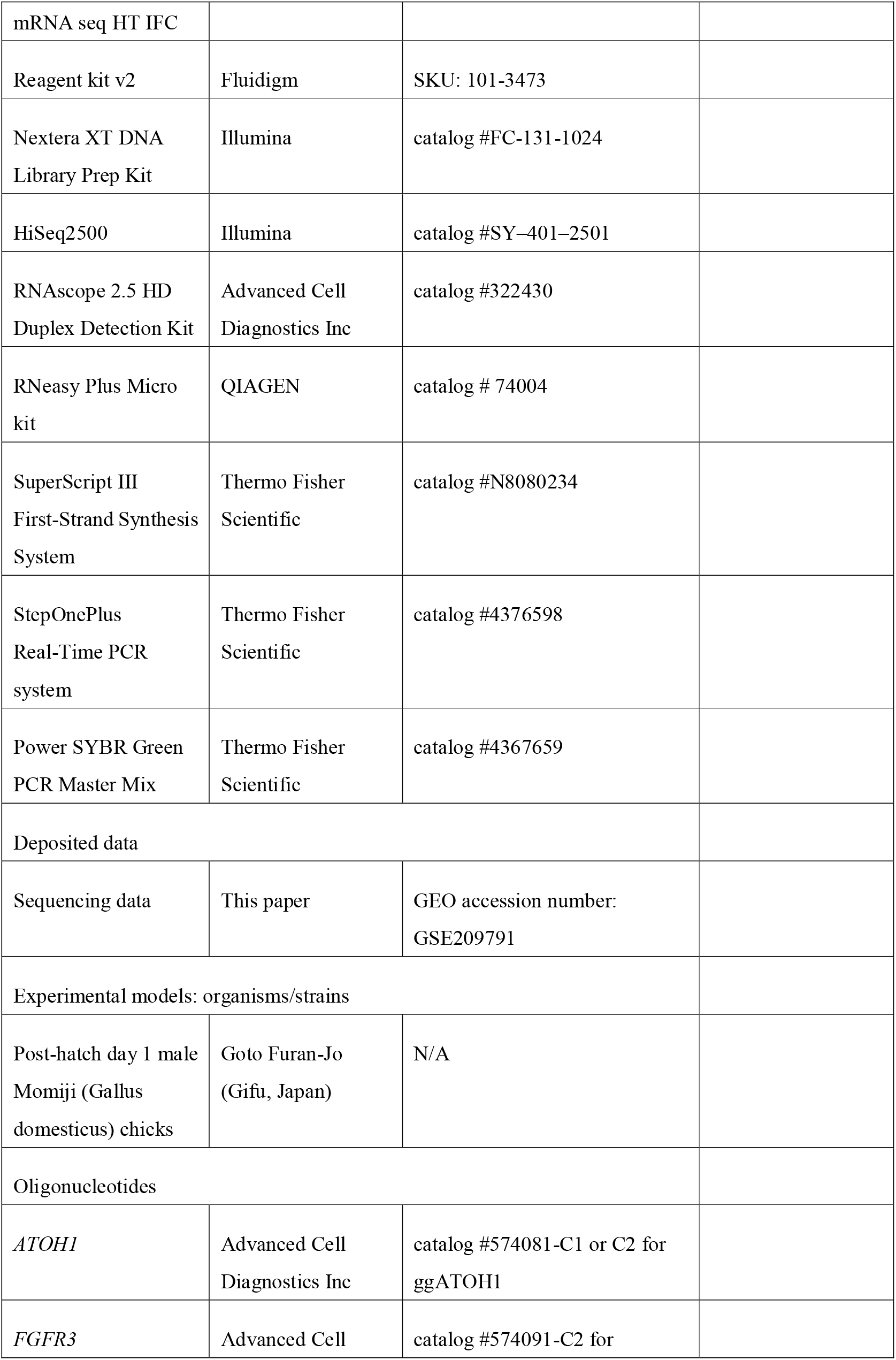
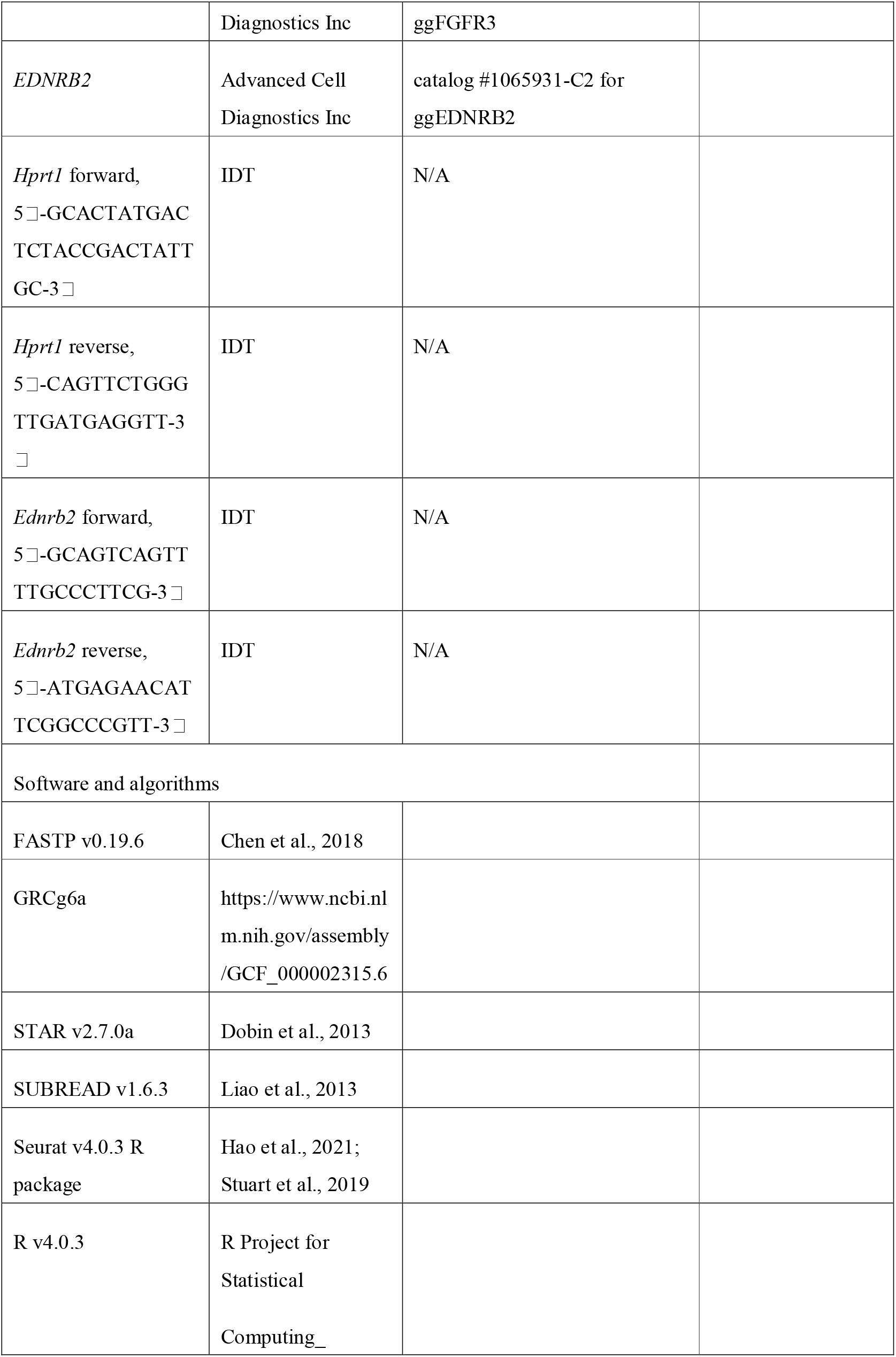
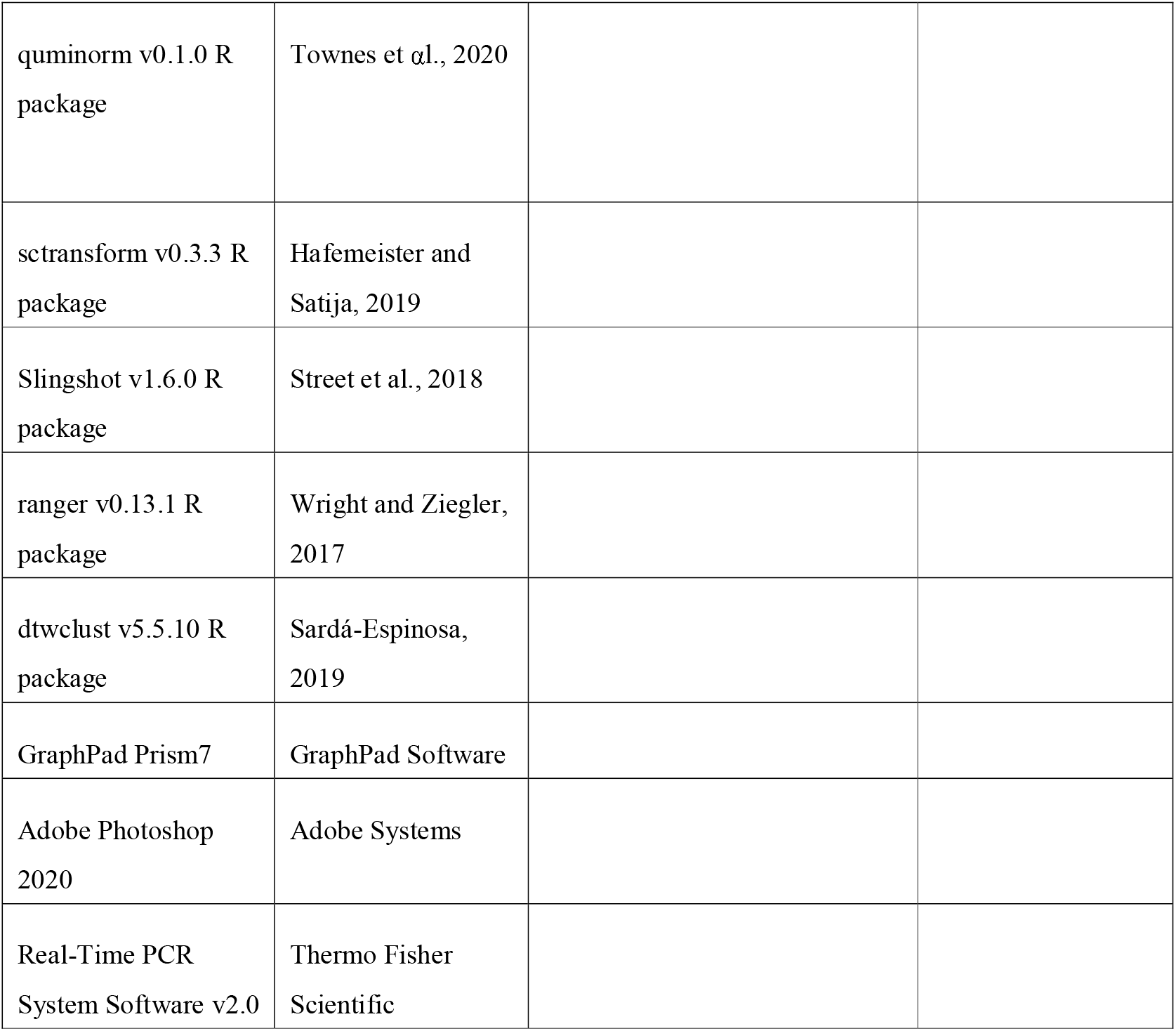

### Explant culture of chick basilar papillae

Post-hatch day 1 (P1) male Momiji (Gallus domesticus) chicks were purchased from Goto Furan-Jo (Gifu, Japan) and placed for less than 3 h in the transport box in the experimental room, which was maintained at approximately 23°C. All animal procedures were performed following the National Institutes of Health (NIH; Bethesda, MD, USA) Guide for the Care and Use of Laboratory Animals (NIH Publications No. 8023, revised 1978) and were approved by the Animal Research Committee of Kyoto University Graduate School of Medicine (MedKyo21123).

The basilar papilla (BP) explant culture was performed according to our previous study (Matsunaga et al., 2020). Briefly, the middle ear cavity was opened, followed by the extirpation of the cochlear duct. Cochlear ducts were placed in ice-cold sterile medium 199 (catalog #11150059, Thermo Fisher Scientific, Waltham, MA, USA). The tegmentum vasculosum was carefully removed from the cochlear duct. The remaining tissue containing the BP was cultured in a free-floating manner (submerged in the medium) in 500 µl control medium consisting of Dulbecco’s modified Eagle’s medium with 4.5 g/l glucose (DMEM-high glucose, catalog #08459-35, Nacalai Tesque, Kyoto, Japan) supplemented with 1% fetal bovine serum (FBS, catalog #12389802, Thermo Fisher Scientific) in 48-well plates (Iwaki Co. Limited, Tokyo, Japan) for 24 h. Thereafter, BP specimens were provided for explant cultures. To damage hair cells (HCs), BP explants were maintained with the culture media containing penicillin/streptomycin (SM; 200 penicillin units per ml and 78 µM streptomycin, Sigma–Aldrich, St. Louis, MO, USA, catalog #P4333). After the SM exposure for 48 h, BP was washed out by changing the medium twice, and then cultured in DMEM-high glucose with 1% FBS and penicillin (control medium). The control specimens were incubated in the control media without exposure to SM. Culture media were changed every other day.

### Single cell preparation and RNA-sequencing

The following samples were provided for the single cell RNA sequencing (RNA-seq) analysis: day0_pre (before SM exposure, n = 17), day2_SM (immediately after 48-h SM exposure, n = 16), day6_SM (96 h after SM exposure, n = 15), day2_Ctrl (48-h culture in the control medium, n = 16), day6_Ctrl (144-h culture in the control medium, n = 15) and intact BPs that were harvested immediately after the dissection from the temporal bones (n = 17). For enzymatic dissociation of BPs for single cell RNA-seq, specimens were incubated in prewarmed 500 µg/ml thermolysin (catalog #T7902, Sigma–Aldrich) in medium 199 (catalog #11043023, Thermo Fisher Scientific) at 37[for 30 min. To stop enzymatic dissociation, specimens were transferred into medium 199 supplemented with 5% FBS. Afterwards, the BP, the sensory epithelium was separated from the surrounding tissues under the stereomicroscope. Then, the samples were transferred into an Eppendorf tube by a pipette. After centrifugation (800g x 5 min) at room temperature, the supernatant was removed. Samples were incubated in an enzyme cocktail containing 50% Accutase and 50% TrypLE Select enzyme (1X) (1:1, catalog#12679-54, Nacalai Tesque, catalog# 12563011, Thermo Fisher Scientific) for 7 min at 37[. After gently pipetting 10-20 times, samples were incubated for additional 4 min at 37[. After inactivation of enzymatic activity with medium 199 supplemented with 5% FBS, further mechanical dissociation by pipetting was performed. Dissociated cells were collected after centrifugation (800g x 5 min) and resuspended in medium 199 supplemented with 5% FBS. We sampled 1 µl of a cell suspension and quantified the number of living cells by a TC20 automated cell counter (Bio-Rad Laboratories, Hercules, CA USA). Finally, the cell concentration was adjusted to 500 cells per µl.

Capture of the single cells and cDNA synthesis were conducted using C1 Single-Cell mRNA seq HT IFC for medium cell size (10-17 µm) and Reagent kit v2 (Fluidigm, San Francisco, CA, USA), according to manufacturer’s instruction. Briefly, 12,000 cells of each sample were loaded onto HT IFC, and 400 cells were captured in the IFC. Sequencing libraries were prepared by transposase-assisted tagmentation and enrichment of 3’end of genes using Nextera XT DNA Library Prep Kit (Illumina, San Diego, CA, USA). The libraries were sequenced on the HiSeq2500 for 25 and 75 cycles paired-end reads. The raw sequencing data in bcl format were converted into FASTQ files using the Illumina bcl2fastq2 software.

### Data preprocessing

We applied C1 mRNA Seq HT Demultiplex Script v2.0 (Fluidigm) to demultiplex the FASTQ files. We performed quality check and trimming using FASTP v0.19.6 (Chen et al., 2018) with default settings otherwise additional polyG and polyX tail trimming. Trimmed reads were mapped to the chicken reference genome GRCg6a (https://www.ncbi.nlm.nih.gov/assembly/GCF_000002315.6) using STAR v2.7.0a (Dobin et al., 2013) with default settings. The generated BAM files were transformed into gene expression matrix of 2,400 cells using featureCounts function from SUBREAD v1.6.3 package (Liao et al., 2013). We performed a downstream analysis using Seurat 4.0.3 library (Hao et al., 2021; Stuart et al., 2019) on R 4.0.3 (R Core Team (2020). R: A language and environment for statistical computing. R Foundation for Statistical Computing, Vienna, Austria. URL https://www.R-project.org/.). After removing the cells of which RNA count was less than 20,000, detected genes was less than 1,000, detected gene was more than 4,000, or percentage of mitochondrial genes was more than 12.5 %, 1,054 cells were analyzed. The dataset was normalized using transcripts per million (Wagner et al., 2012), quasi UMI normalization (Townes et al, 2020) and R package sctransform, which included the selection of variable genes (Hafemeister and Satija, 2019). We performed principal component analysis (PCA) to reduce dimensionality on the normalized data matrix. For PCA, we used the top 3,000 most highly variable genes and kept the first 10 principal components (PCs). For 2-dimentional visualization, PC embedding was passed into the uniform manifold approximation and projection (UMAP) (McInnes et al., 2018). The parameters for UMAP were as follows: min dist = 0.01, n (the number of neighbors) = 30. We applied the Louvain clustering algorithm for community detection. The Seurat FindMarkers function was used to explore marker genes for the clusters (Wilcoxon rank-sum test). The clusters were annotated based on established marker genes. During analyses of differentially expressed genes, we set a log-fold-change threshold > 0.5 and FDR < 0.05. *P*-value was adjusted by Bonferroni correction. Reconstruction of the pseudotime trajectory was performed by Slingshot v1.6.0 (Street et al., 2018). In order to find genes that are differentially expressed along the pseudotime, we built a random forest regression model that predicts the value of the pseudotime from the expression value of each gene. We built the model using R ranger package (Wright and Ziegler, 2017. The parameters for the regression model were as follows: mtry (randomly selected predictors) = 800, trees (number of trees) = 1,000, min_n (minimal node seize) = 15. Using this model, we calculated the importance of the genes at the pseudo-time and selected 500 most important genes. We classified the 500 selected genes into six clusters using dtwclust package (Sardá-Espinosa, 2019). We performed time series clustering based on Euclidean distances. Volcano plots were made in GraphPad Prism7 (GraphPad Software, San Diego, CA, USA).

### Immunohistochemistry

Immunostaining was carried out on whole mount preparations or frozen sections of the chick cochlea duct. After the fixation with 4% paraformaldehyde (PFA) in phosphate-buffered saline (PBS) for 15 min at room temperature, PBS wash was done for two times. The dehydration was performed for the frozen section from 15% sucrose with 0.2mM ethylenediaminetetraacetic acid (EDTA) in 1×PBS at 4 °C overnight to 30% sucrose with 0.2mM EDTA next night. Specimens were embedded in Tissue-Tek O.C.T. Compound (Sakura Fine technical Co., Ltd., Tokyo, Japan) and then, cryostat-cut sections (10 µm) were mounted directly onto MAS-coated slides (Matsunami Glass Ind. Ltd, Osaka, Japan). The ensemble of sections collected from 2-3 BPs was distributed through almost 20 slides (7-9 sections per slide) for each experiment. And each slide contains a reliable representation of the whole length of the BP in a serial manner (first tissue section on slide numbered 1 and the following tissue section on slide number 2 and so on) which provided uniformity of treatment for cross-sections throughout the cochlear length.

Both whole-mount samples and section samples were incubated in blocking solution (1% bovine serum albumin and 5% normal goat serum in PBS) for 30 min at room temperature. The samples were incubated with primary antibodies in a blocking solution overnight at 4 °C, followed by incubation with the corresponding secondary antibodies for 1 h at room temperature. Secondary antibodies: Alexa Fluor 546 goat anti-mouse (1:1000, Thermo Fisher, A11003); Alexa Fluor 568 goat anti-mouse (1:1000, Thermo Fisher, A11004); Alexa Fluor 488 goat anti-rabbit (1:1000, Thermo Fisher, A11034); Alexa Fluor 546 goat anti-rabbit (1:1000, Thermo Fisher, A11010). Nuclear staining was done by 4’,6-diamidino-2-phenylindole (DAPI, catalog # D1306, Thermo Fisher Scientific). After several washes in 1xPBS, samples were mounted using FluoromountG (catalog # 00-4958-02, Southern Biotech, Birmingham, AL, USA). All fluorescence images were obtained with a TCS-SPE confocal microscope (equipped with 40×/1.15 OIL CS. objective, Leica Microsystems, Wetzlar, Germany). As for whole mount samples, each image taken by the BX50 microscope (Olympus, Tokyo, Japan) was identified at first by immunostaining for MYO7A and nuclear staining with DAPI, and the total BP length between the proximal portion and the distal portion was calculated by ImageJ software (NIH, https://imagej.nih.gov/ij/). Around 40% of the distal end of each BP was used for quantitative analyses in whole-mount samples. Optical sections in the xy-field (“z-sections”) were imaged and recorded at 4-μm intervals with the span adjusted to include the HC layer and SC layer in the xy-field of view. As for transverse sections, 3 sections for each image were z-stack projection. Image processing for figures was accomplished in Adobe Photoshop 2020 (Adobe Systems, San Jose, CA, USA). Images presented are representative of three independent experiments. We counted numbers of MYO7A-expressing cells with DAPI nuclear staining as hair cell numbers and those of MYO7A and SOX2 co-expressing cells as immature hair cell numbers. Welch t-test for the two groups was calculated in GraphPad Prism 7.

### *In situ* hybridization

The preparation of the frozen-section sample for the *in situ* hybridization (ISH) was almost the same as immunostaining protocols except for the longer fixation (at 4 °C for 3-4 h), higher EDTA concentration (333mM) in 30% sucrose and thicker sections (14 µm). All ISH experiments were carried out with the RNAscope 2.5 HD Duplex Detection Kit (Advanced Cell Diagnostics Inc., Hayward, CA, USA) on frozen section samples according to the manufacture’s instruction. Briefly, slides were baked at 60 °C for 15 min and washed with 1xPBS. The specimens were then post-fixed in pre-chilled 4% PFA for 15 min, washed in 2 changes of double distilled water (DDW) for 1 min each before dehydration through 50%, 70%, 100% and 100% ethanol for 5 min each. The slides were air-dried for 5 min before hydrogen peroxide treatment for 10 min, washed with DDW for a brief time twice. Slides were boiled in the target retrieval solution for 3 min and washed with DDW and then changed for 100% ethanol. Slides were dried up again with baking at 60 °C for 10 min and then allowed to put into the Protease III Solution at 40 °C for 5 min, and the hybridization with the pre-warmed probe was performed at 40 °C for 120 min. After washing with wash buffer and 5xSSC solution overnight, the specimens were stained according to the manufacturer’s instructions. On each individual section, three target mRNAs were examined: *ATOH1* (#574081-C1 or C2 for ggATOH1, Advanced Cell Diagnostics Inc.), *FGFR3* (574091-C2 for ggFGFR3, Advanced Cell Diagnostics Inc.), and *EDNRB2* (1065931-C2 for ggEDNRB2, Advanced Cell Diagnostics Inc.). Both positive and negative control probes (#453961-C2 for ggUBC [ubiquitin C] or #320751, Advanced Cell Diagnostics Inc.) were also labeled. The duplex negative control probe (#320751, Advanced Cell Diagnostics Inc.) was used on one section per slide for control purposes. Probes in channel 2 were labeled with AP enzyme and a red substrate. DAPI was used to mark cell nuclei. Slides were then imaged using a BX50 microscope to take 20x, 40x bright-field and fluorescence with DAPI filter from more than 3 BP samples for each time point, and a representative image was cited in the figures. Image processing for figures was accomplished in Adobe Photoshop 2020 (Adobe Systems). Images presented are representative of three independent experiments.

### Quantitative real-time polymerase chain reaction

Total RNA for each sample was extracted pooled 10 BPs using the RNeasy Plus Micro kit (catalog # 74004, QIAGEN, Venlo, Netherlands) according to the manufacturer’s protocol. DNase I treatment was performed using spin columns. RNA was reverse-transcribed using the SuperScript III First-Strand Synthesis System (catalog #N8080234, Thermo Fisher Scientific, Waltham, MA, USA). Quantitative real-time polymerase chain reaction (qPCR) was performed using a StepOnePlus Real-Time PCR system (Thermo Fisher Scientific, Waltham, MA, USA). cDNA was amplified using the Power SYBR Green PCR Master Mix (catalog #4367659, Thermo Fisher Scientific, Waltham, MA, USA). The experiment was performed in triplicate. Target gene expression was normalized to hypoxanthine phosphoribosyltransferase 1 (Hprt1, Hassanpour et al., 2019). cDNA from the brain tissue of post-hatched 1-day-old chickens was used to generate standard curves for each gene. Relative quantification was performed using Real-Time PCR System Software v2.0, with the 2−ΔΔCT method (Thermo Fisher Scientific, Waltham, MA, USA). The following primers were used:

*Hprt1* forward, 5′-GCACTATGACTCTACCGACTATTGC-3′

*Hprt1* reverse, 5′-CAGTTCTGGGTTGATGAGGTT-3′

*Ednrb2* forward, 5□-GCAGTCAGTTTTGCCCTTCG-3□

*Ednrb2* reverse, 5□-ATGAGAACATTCGGCCCGTT-3□

### Pharmacological inhibition of transforming growth factor **β** type 1 receptors

Chick BP explants were cultured with SM (78 µM) alone (day2_SM) or SM and a transforming growth factor β type 1 receptor inhibitor, SB505124 (catalog #S4696-5MG, Sigma-Aldrich) at a concentration of 2μM (day2_SM+SB) for 48 h. To validate *EDNRB2* expression, we harvested BP explants after 48-h culture and performed qPCR and ISH. For qPCR assessment related to Figure 5B, we used 10 to 11BPs for each experimental group per experiment. BPs before SM exposure were used as controls (day0_pre). For quantification of *EDNRB2*-expressing cells related to Figure 5C, we counted *EDNRB2*-expressing cells with DAPI staining in three transverse sections of BPs from the distal (0-30% from the distal end), mid (30-70%) and proximal (70%-the proximal end) portions in each sample. The sum of three sections was defined as the number of *EDNRB2*-expressing cells for each sample. To examine effects of pharmacological inhibition of transforming growth factor β type 1 receptor on hair cell regeneration, we used BP explants that were cultured with SM or SM+SB505124 for 48 h followed additional 96-h culture in control media, and HC count was done in transverse sections using immunohistochemistry for MYO7A and nuclear staining with DAPI. Welch t-test for the two groups was calculated in GraphPad Prism 7.

## Acknowledgements

We thank J.S. Stone and her lab members (Univ. of Washington) for technical advice regarding explant cultures of chick basilar papillae, S. Heller (Stanford Univ.) for practical advice regarding tissue preparation for *in situ* hybridization of chick basilar papillae, A. Watanabe and S. Sakamoto (Kyoto Univ.) for performing single cell-RNA sequencing, and S. Suzuki, C. Long and H. Doods (Boehringer Ingelheim Pharma GmbH) for discussions regarding bioinformatics analyses. The study was partly supported by KAKENHI (Grants-in-Aid for Scientific Research, 20K09708 to T.N., Grant-in-Aid for Young Scientists [Start-up], 21K20964 to M.M., Grant-in-Aid for Young Scientists [B], 22K16899 to M.M.) from the Japan Society for the Promotion of Science, by Start-up FY2021 to M.M. from Kyoto University Research Administration Office, by AMED (Japan Agency for Medical Research and Development) under Grant (20lm0203013j0002) to T.O., and by Boehringer Ingelheim Pharma GmbH to T.N..

## Author Contributions

Conceptualization, M.M. and T.N.; methodology, R.Y., T.K., M.M. and T.N.; formal analysis, M.M., R.Y., T.K., and T.N.; investigation, M.M., R.Y., T.K., and H.O.; resources, M.M., T.O, N.Y., K.O., and T.N.; writing – original draft, M.M., R.Y., T.K., and T.N.; writing – review & editing, T.O, N.Y., and T.N.; visualization, M.M., R.Y. and T.N.; supervision, N.Y., K.O., and T.N.; funding acquisition, M.M., T.O., and T.N.

## Legends for supplementary figures and tables

**Figure S1.**
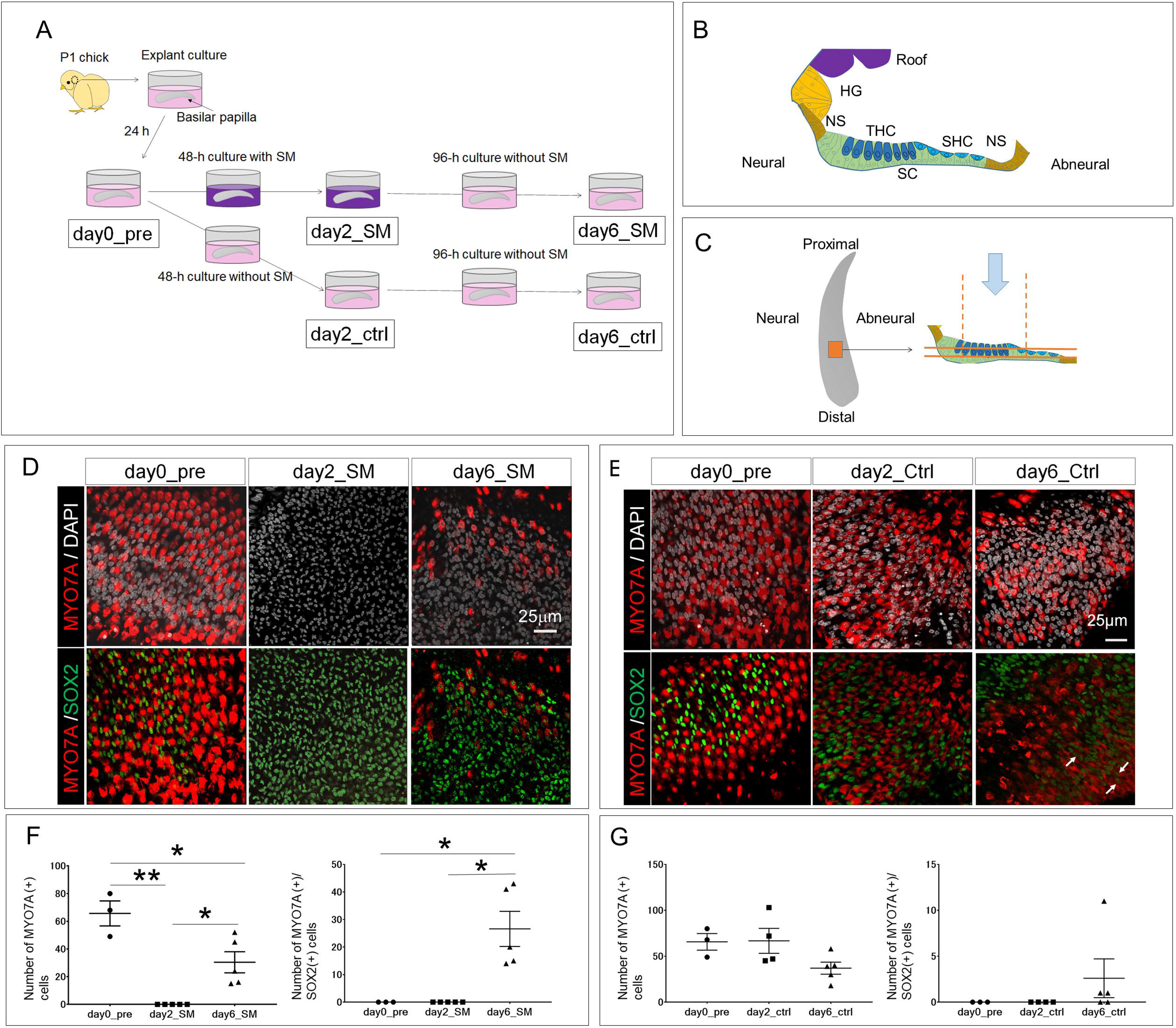
Hair cell regeneration occurs in explant cultures of chick basilar papillae. (A) Experimental design and experimental groups. Specimens that were collected before explant culture (day0_pre, n=3), those after 48-h exposure to streptomycin (SM) (day2_SM, n=5), those cultured for additional 96 h after SM exposure (day6_SM, n=5) and those cultured 48 h (day2_Ctrl, n=4) or 144 h (day6_Ctrl, n=5) without SM exposure were used. (B) Schematic drawing of a transversal section of the basilar papilla. The sensory epithelium consists of hair cells (tall [THC] and short hair cell [SHC]) and supporting cells (SC). Non-sensory cells (NS) are adjacent to the neural and abneural edge of the sensory epithelium. A subtype of non-sensory cells, homogene cells (HG) are present in the neural side of the cochlear duct and connect with the cochlear roof cells (Roof). (C) Schematic drawing of the observation area in the surface of a chick basilar papilla and two layers for cell counting. (D, E) Confocal images of the 40% area from the distal end of basilar papilla surfaces that are stained with immunohistochemistry for MYO7A (red) and nuclear staining with DAPI (grey) or stained with immunohistochemistry for MYO7A (red) and SOX2 (green) before and after SM exposure (D) and without SM exposure (E). Arrows indicate MYO7A and SOX2-positive cells. Scale bars represent 25 μm. (F, G) Hair cells labeled with MYO7A are totally lost after 48-h SM exposure (day2_SM) and a significant recovery in numbers of MYO7A-positive cells is observed in day6_SM specimens (F). A significant increase of cells co-stained with MYO7A and SOX2 is observed in day6_SM specimens (F). In specimens that were cultured without SM exposure, no significant differences in numbers of MYO7A-positive cells or MYO7A and SOX2-positive cells are found between experimental groups (G). Error bars represent SEM. One-way ANOVA with Tukey’s correction was used to calculate *P* values. \**P* < 0.01, \*\**P* < 0.001.

**Figure S2.**
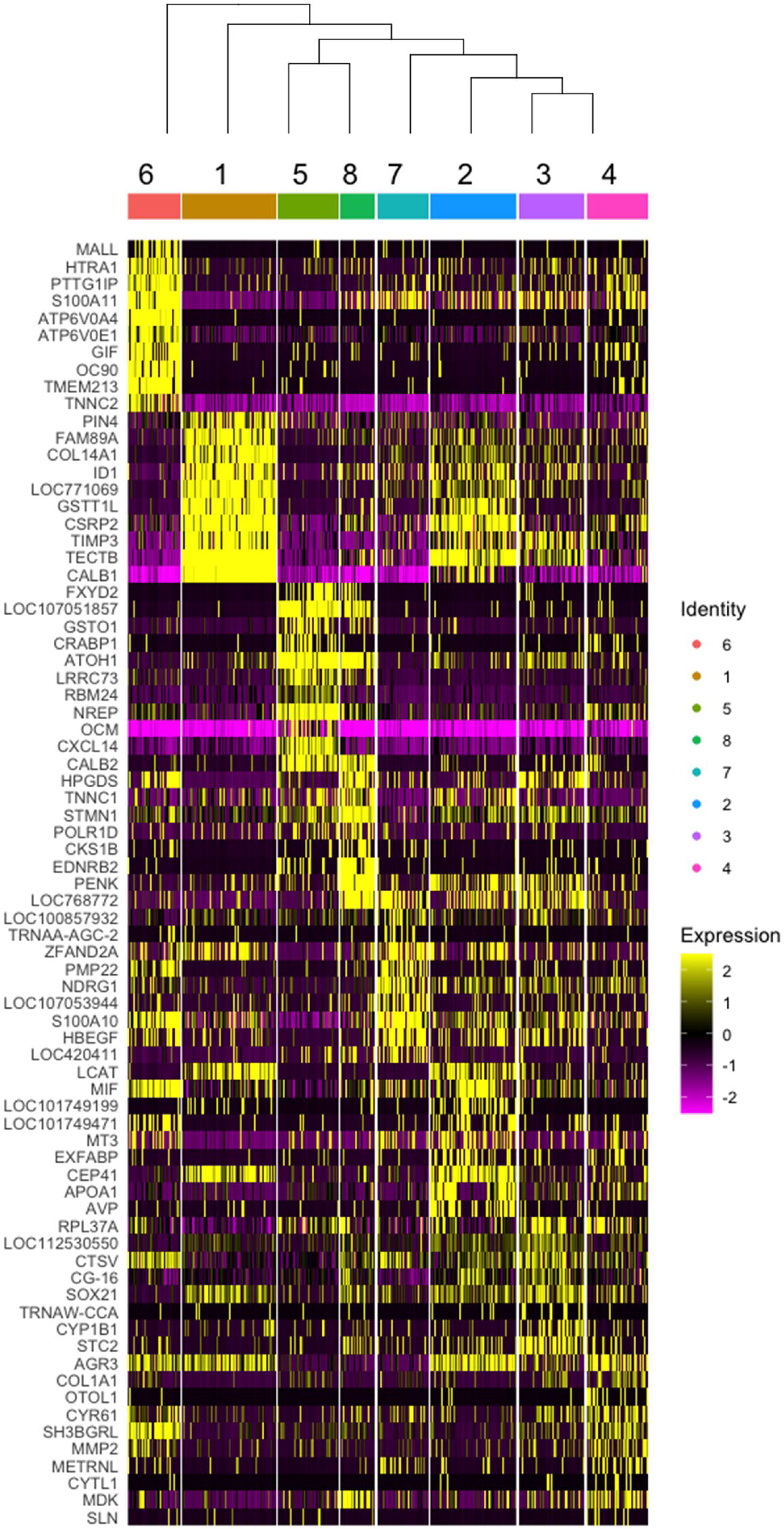
Heatmap of representative genes for SC-clusters and hierarchy of SC-clusters.

**Figure S3.**
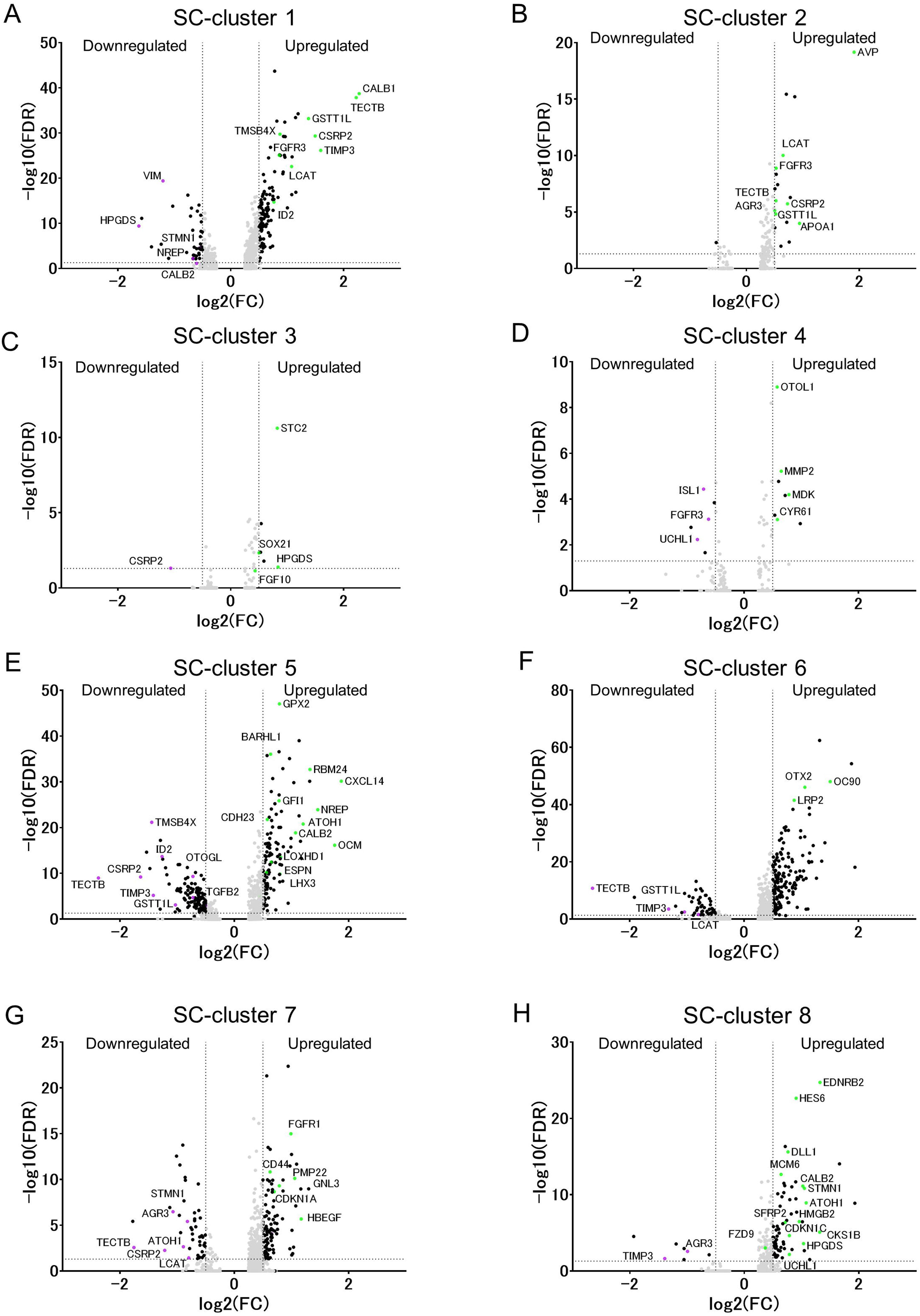
Volcano plots showing downregulated and upregulated genes in each SC-clusters compared with the other SC-clusters (FDR<0.05, log_2_[fold change]>0.5 or <-0.5).

**Figure S4.**
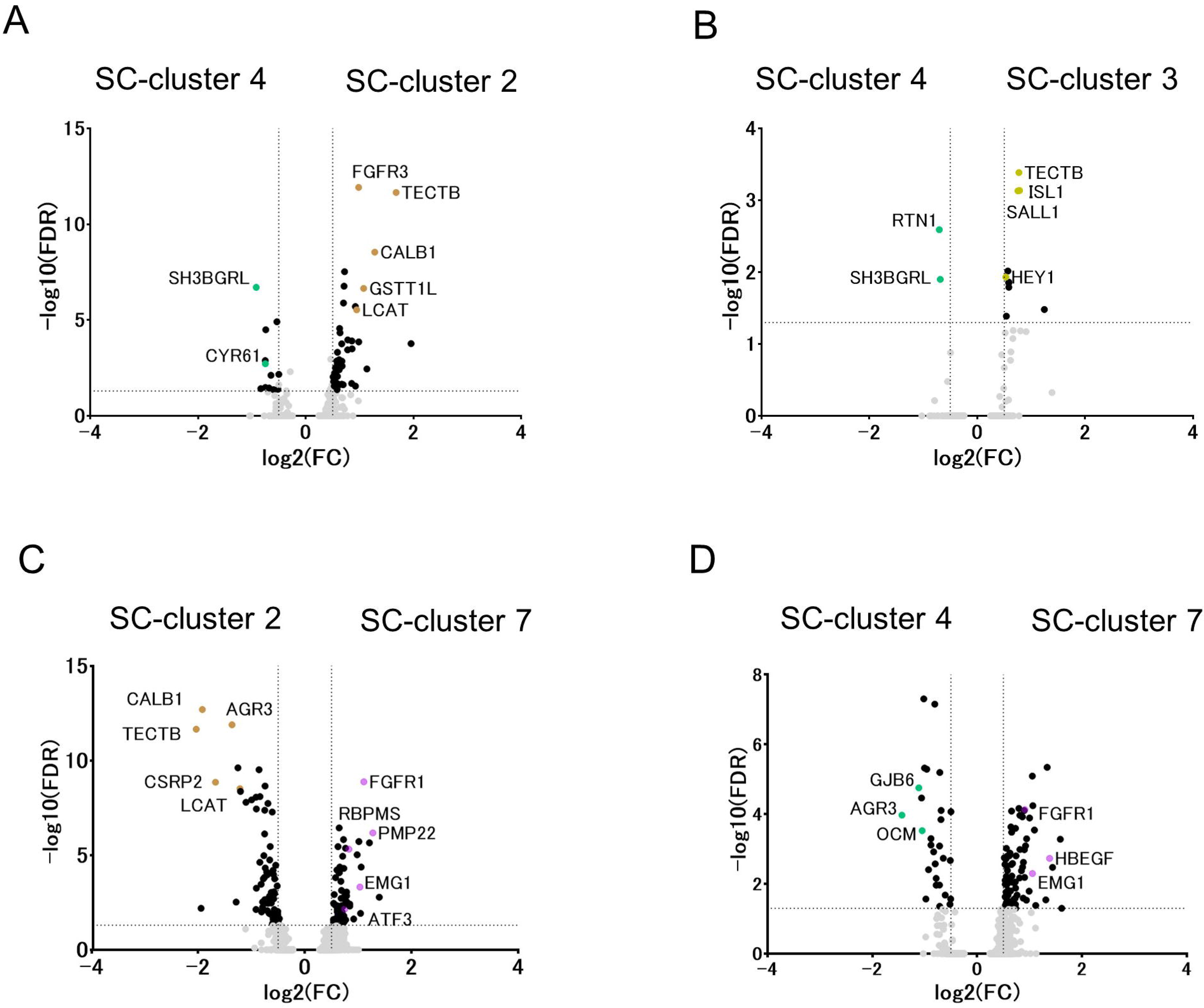
Volcano plots showing highly expressed genes (FDR<0.05, log-fold-change threshold > 0.5) in comparison between SC-cluster 4 and 2 (A), SC-cluster 4 and 3 (B), SC-cluster 2 and 7 (C), and SC-cluster 4 and 7 (D).

**Figure S5.**
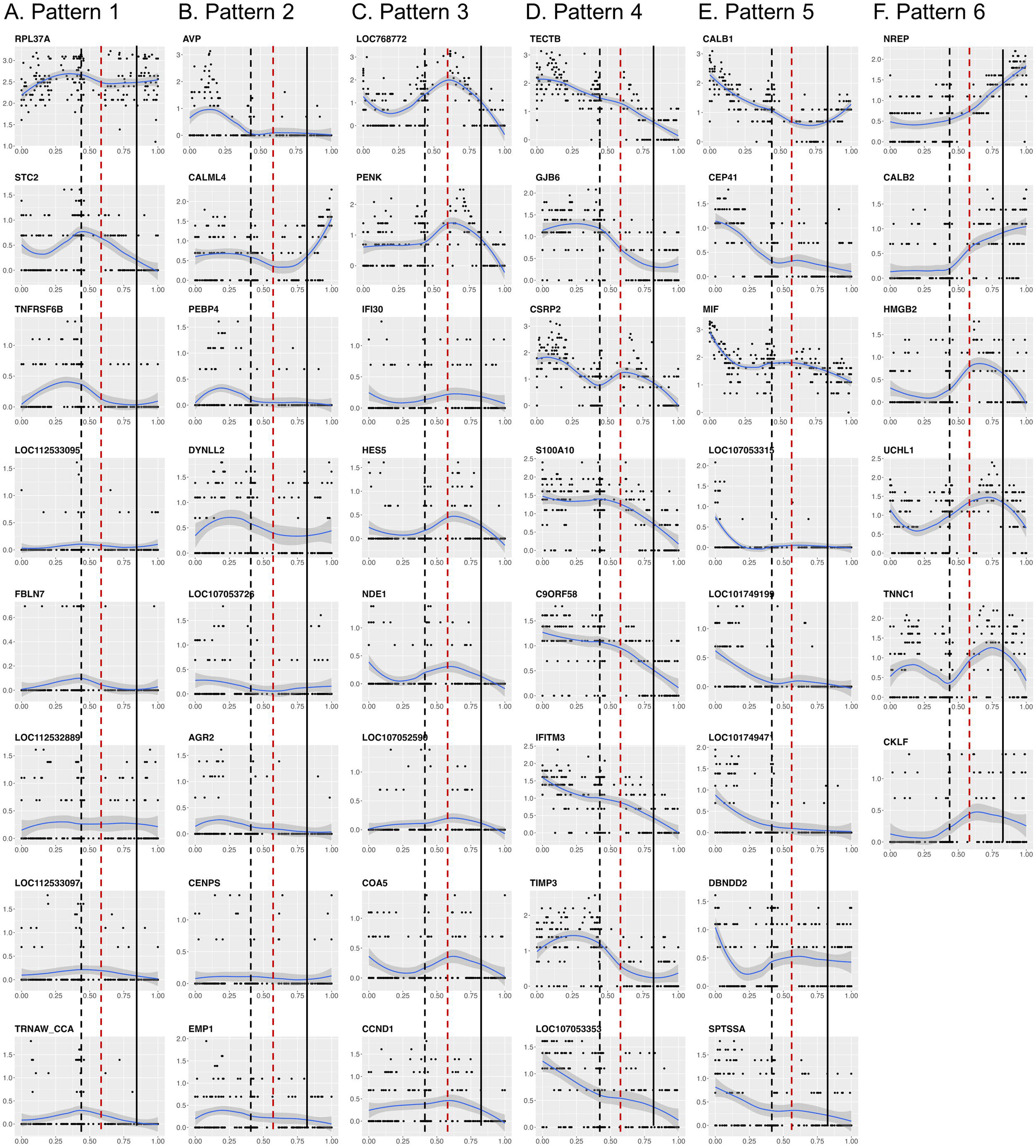
Normalized expression values of genes listing in Figures 4D-I as top 10 genes along a pseudotime line for pattern1 (A), 2 (B), 3 (C), 4 (D), 5 (E) and 6 (F). Black dotted, red dotted and black lines indicate the timing for *EDNRB2* induction, for the induction of hair cell differentiation genes and for *ATOH1* downregulation, respectively.

**Figure S6.**
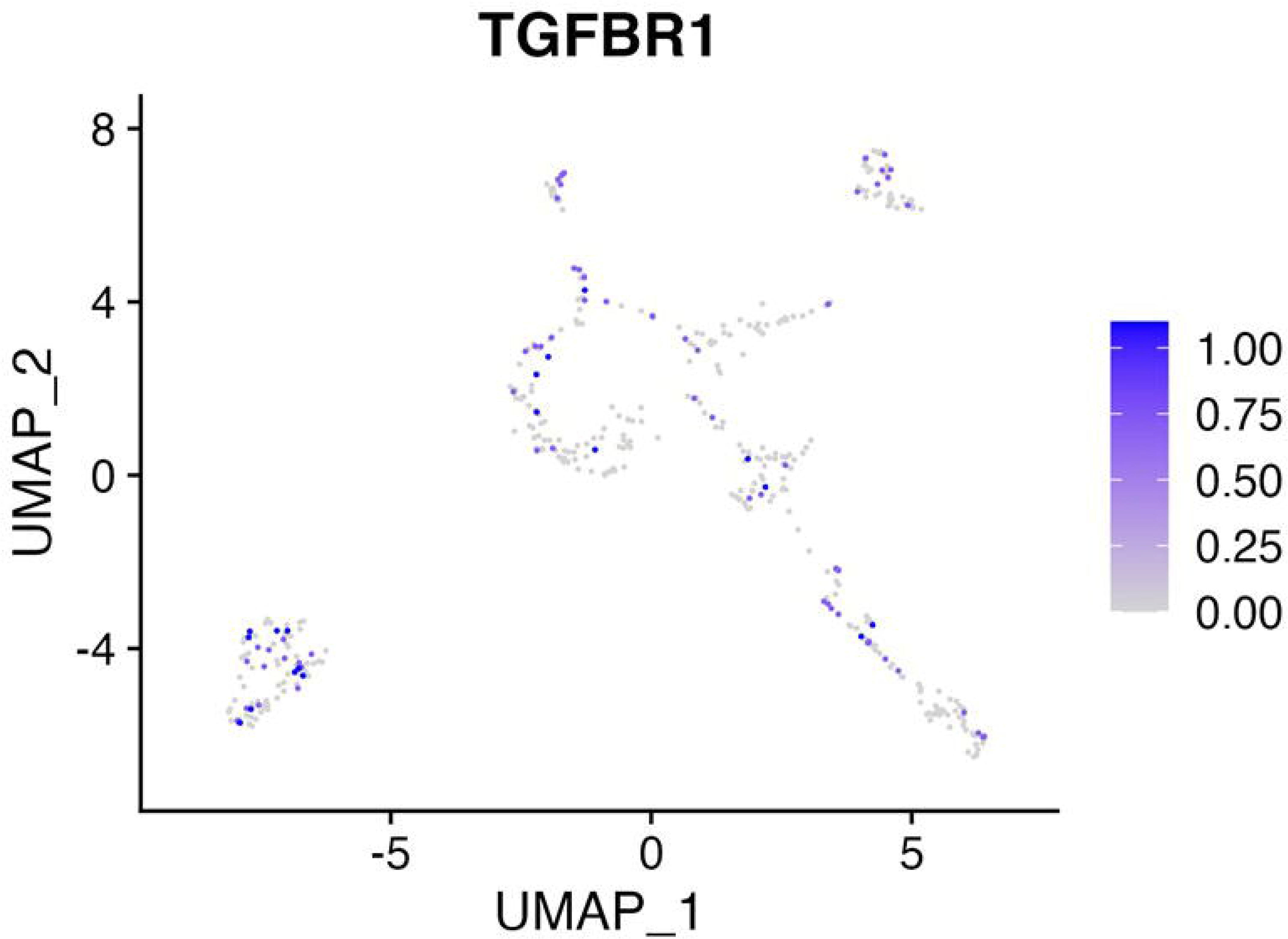
Uniform manifold approximation and projection plots for TGFBR1 in SC-clusters.

**Figure S7.**
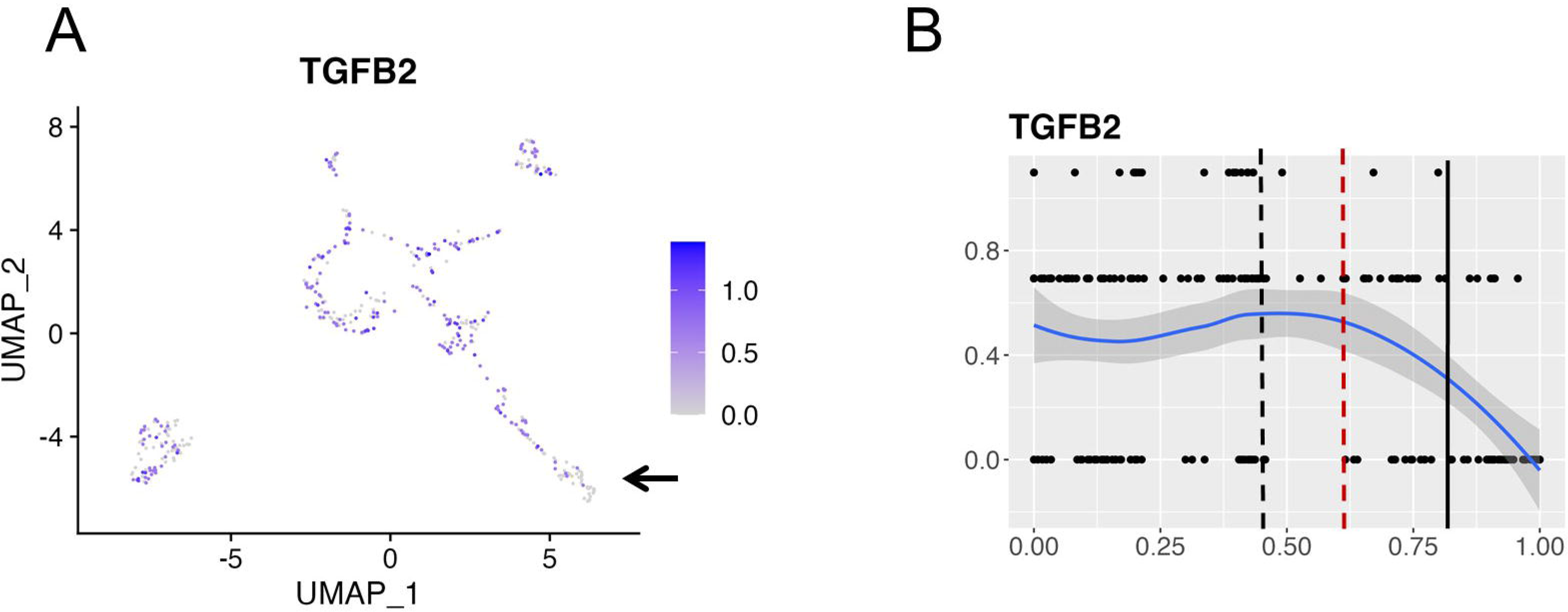
Uniform manifold approximation and projection plots (A) and pseudotime trajectory for TGFB2 (B). An arrow (A) indicates a cluster showing low expression of *TGFB2*. Black dotted, red dotted and black lines indicate the timing for *EDNRB2* induction, for the induction of hair cell differentiation genes and for *ATOH1* downregulation, respectively (B).

**Table S1.** Dataset of clusters consisting of supporting cells and neighboring non-sensory cells for differentially expressed genes corresponding to Figure S3.

**Table S2.** Dataset for differentially expressed genes between SC-cluster 5 and 8, between SC-cluster 8 and 3, between SC-cluster 8 and 2 and between SC-cluster 2 and 3 corresponding to Figure 3.

**Table S3.** Dataset for differentially expressed genes between SC-cluster 2 and 4, between SC-cluster 3 and 4, between SC-cluster 2 and 7 and between SC-cluster 4 and 7 corresponding to Figure S4.

**Table S4.** Dataset of 500 differentially expressed genes along a pseudotime trajectory by fitting a random forest regression model corresponding to Figures 4 and S5.

